# Proteostasis Landscapes of Cystic Fibrosis Variants Reveals Drug Response Vulnerability

**DOI:** 10.1101/2024.07.10.602964

**Authors:** Eli Fritz McDonald, Minsoo Kim, John A. Olson, Jens Meiler, Lars Plate

## Abstract

Cystic Fibrosis (CF) is a lethal genetic disorder caused by variants in CF transmembrane conductance regulator (CFTR). Many variants are treatable with correctors, which enhance the folding and trafficking of CFTR. However, approximately 3% of persons with CF harbor poorly responsive variants. Here, we used affinity purification mass spectrometry proteomics to profile the protein homeostasis (proteostasis) changes of CFTR variants during correction to assess modulated interactions with protein folding and maturation pathways. Responsive variant interactions converged on similar proteostasis pathways during correction. In contrast, poorly responsive variants subtly diverged, revealing a partial restoration of protein quality control surveillance and partial correction. Computational structural modeling showed that corrector VX-445 failed to confer enough NBD1 stability to poor responders. NBD1 secondary stabilizing mutations rescued poorly responsive variants, revealing structural vulnerabilities in NBD1 required for treating poor responders. Our study provides a framework for discerning the underlying protein quality control and structural defects of CFTR variants not reached with existing drugs to expand therapeutics to all susceptible CFTR variants.

**SIGNIFICANCE STATEMENT:** Cystic Fibrosis (CF) is a lethal genetic disease with variants leading to misfolding of an anion channel protein. Enhancing productive channel folding using a novel class of small molecules called *correctors* has emerged as the current CF treatment paradigm. However, correctors fail to reach all patient variants. Using high throughput interactomics, Rosetta simulations, and biochemical trafficking assays, this study demonstrates poorly responsive CF variants experience diverse misfolding pathways caused by structural defects in the core of a nucleotide-binding domain. Stabilizing secondary mutations in this domain rescues poorly responsive variants, paving the way for mechanistic-based therapeutic development for untreatable CF variants and future protein misfolding corrector drugs.

**COMPETING INTERESTS:** The authors declare no competing interests.

## INTRODUCTION

Cystic Fibrosis is a lethal genetic disease caused by mutations in Cystic Fibrosis Transmembrane Conductance Regulator (CFTR), an epithelial anion channel. Mutant CFTR leads to a lack of anion transport in the lung epithelia, dysfunctional cilia transport of mucus, mucus build-up, infection, and eventually death. CFTR variants are broadly categorized as Class-I (non-sense variants), Class-II (misfolding-prone variants), or Class-III (mis-gating variants)^1^. The current treatment strategy for CF involves small-molecule therapeutics that either increase Class-I variant translation efficiency (read-through agents), elevate Class-III channel activity (potentiators), or bolster Class-II folding stability (correctors)^1,2^. Correctors were originally developed to treat the most common CFTR variant, the deletion of phenylalanine at position 508 (F508del). Beyond F508del, the FDA has approved the combination therapy of two correctors (VX-661 & VX-445) and one potentiator (VX-770) for 183 other CF-causing variants^2,3^. CFTR variant response is defined by surface trafficking, plasma membrane (PM) expression, and functional rescue of anion conductance. Treatable variants respond differently to mechanistically distinct correctors, demonstrating *selective response* and creating a need for rational combinatorial therapies guided by variant-specific drug response profiles^2^. Despite the revolution in CF treatments, ∼10% of persons with CF remain untreated^4^. Among the total CF population, ∼3% harbor poorly responsive Class-II variants potentially targetable with correctors. However, sparse biochemical research on mechanisms of response and non-response has hindered progress toward treating poorly responsive variants.

CFTR is an ABC transporter-type protein comprised of two nucleotide-binding domains (NBDs), two transmembrane domains (TMDs), and an unstructured R domain (RD)^5^. Correctors are classified by their capacity to stabilize a domain or a domain interface. Type-I corrector VX-661 binds TMD1^6^ and stabilizes the TMD/NBD1 interface^7^. By contrast, Type-III corrector VX-445 binds to the interface of TMD2 and the N-terminal lasso motif^8^ to allosterically stabilize NBD1^7,9^. Type-II correctors stabilize NBD2, although none are FDA-approved to date^7^.

A previous deep mutational scan (DMS) of 129 CFTR variants revealed selectively responsive variants proximate to the VX-445 binding pocket, implying proximity drives selectivity^10^.

Protein homeostasis (proteostasis) is also important for selective response. Misfolding variants continually cycle between proteostasis factors, which direct CFTR to degradation, leading to low steady-state CFTR levels^2,11,12^. We developed a multiplexed, quantitative interactomics platform for measuring changing CFTR interactions with protein homeostasis machinery or *proteostasis landscapes*^13,14^. Our platform demonstrated a role for proteostasis in P67L CFTR response to VX-809^13^ (a VX-661 analog) and VX-445^14^. Thus, correctors modulate CFTR proteostasis in addition to stabilizing proximate structural defects.

The previous DMS study also revealed several poorly responsive variants, such as V520F, L558S, A559T, R560T, A561E, Y569D, and R1066C^15^. Prior research has focused little on these variants because of their low allele frequency (**Supplemental Table S1**). However, a few previous *in vitro* studies showed no or mild response^9,16–19^. Importantly, small molecules and stabilizing mutations had varying effects on poor responders, indicating different capacities for correction. Thus, we sought to understand the proteostatic and structural deficiencies of poorly responsive variants to reveal commonalities, disparities, and vulnerabilities to correction.

Here, we used quantitative multiplexed interactomics to compare the proteostasis landscapes of five VX-445 selectively responsive variants – L165S, Y1032C, T1036N, H1054D, and R1066H - and seven poorly responsive variants – V520F, L558S, A559T, R560T, A561E, Y569D, and R1066C. Responsive variants experience divergent basal interactions with proteostasis pathways, but these proteostasis landscapes converge under VX-445 treatment. By contrast, poorly responsive variants showed subtly different proteostasis changes during dual corrector treatment. Proteostasis landscape remodeling during correction split the poorly responsive variants into *productive* and *trapped* proteostasis variants. Computational structural simulations in Rosetta showed that VX-445 fails to confer enough NBD1 stability to the trapped proteostasis variants. We increased NBD1 stability with secondary mutations, such as I539T, to rescue poorly responsive variants A559T, R560T, and Y569D in combination with correctors. This suggests that increased NBD1 stability may be required to treat poorly responsive variants, paving a path forward for persons with CF without therapeutic options. Thus, vulnerabilities amongst poorly responsive CFTR variants. This may guide future studies to characterize drug susceptible variants for genetic diseases.

## RESULTS

### **I.** Selectively and poorly responsive CFTR variants contact at the NBD1/ICL4 ATPase core interface

While some CFTR variants respond promiscuously to correctors, others demonstrate a selective response to one corrector over another. A previous deep mutational scan (DMS) of CFTR compared VX-661 and VX-445 treatments and revealed L165S, Y1032C, T1036N, H1054D, and R1066H as selectively responsive to VX-445 and P67L, R75Q, Q98R, and V201M as selectively responsive to VX-661^10^ (**Supplemental Figure S1A**). The proximity of selectively responsive variants to the respective VX-445 and VX-661 binding sites led to the paradigm that local intermolecular interactions may drive selective response^6,8^ (**Supplemental Figure S1B**).

The previous DMS study demonstrated poorly responsive variants with <30 % WT cell surface levels (**Figure 1A**). We filtered these previous DMS results against the variants FDA-approved for VX-445 and VX-661 to reveal two hotspots where unapproved, poorly responsive variants occur^3^ (**Figure 1B, Supplemental Figure S1C**). Poorly responsive variants are found at the transition from the alpha-helical subdomain to the ATPase subdomain of NBD1 (**Figure 1C**) (L558S, A559T, R560T, A561E, and Y569D). Additionally, they are found in the intercellular loop 4 (ICL4) region of TMD2 (L1065P and R1066C). Although far in sequence, these variants contact each other in the three-dimensional folded CFTR structure at the NBD1/ICL4 interface (**Figure 1D**).

**Figure 1.**
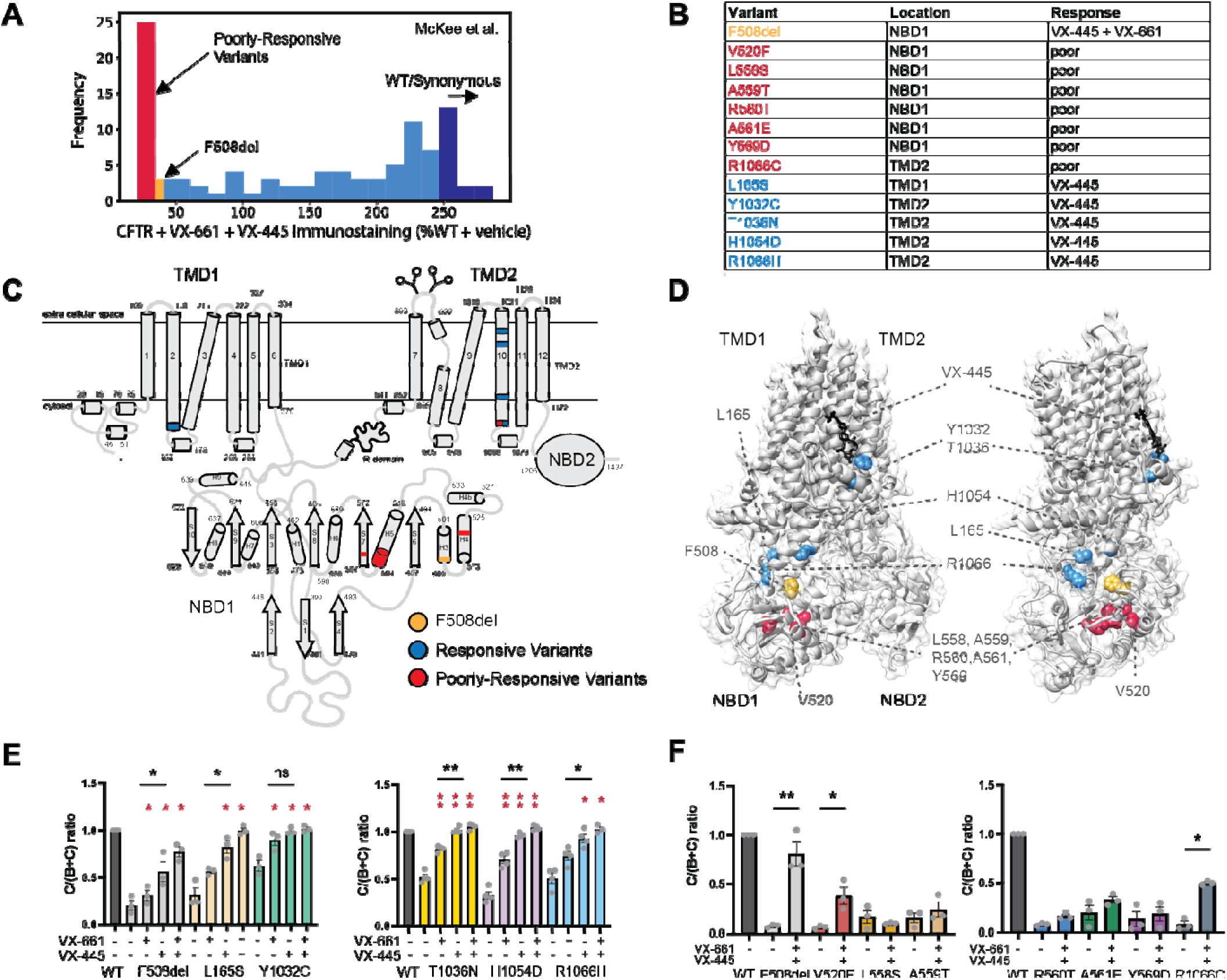
Identification and location of VX-445 selective and poorly responsive CFTR variants. **A.** Histogram of CFTR cell surface immunostaining intensity in percent WT (%WT) under VX-661 and VX-445 dual treatment conditions from McKee et al. 2023 reveals variants that fail to respond to correctors^10^. Poorly responsive variants shown in red, F508del shown in gold, and WT and synonymous variants shown in dark blue. **B.** Table of variants used in this study organized by responsiveness and location in CFTR domains. **C.** Ribbon diagram of the CFTR membrane topology and NBD1 structure with VX-445 selectively responsive variants highlighted in blue and poorly responsive variants highlighted in red. F508 is shown in gold. **D.** CFTR inactive state cryo-EM model (PDB ID 5UAK)^41^ demonstrates VX-445 selectively responsive and poorly responsive variants both cluster in the three-dimensional structure at the ICL4/NBD1 core binding interface. Colored the same as **C.** **E.** Quantification of selectively responsive variants as measured by the C/(C+B) band ratio in response to VX-445 (red asterisks). Difference between VX-445 and VX-661 shown with black asterisks. **F.** Quantification of poorly responsive CFTR variants. Statistical significance was calculated using a parametric ratio paired t-test, and p values are depicted by *< 0.05, **< 0.01, ***< 0.001, and ****< 0.0001. Representative blots for E-F are shown in **Supplemental Figure S1F-G**.

We filtered a larger CFTR variant drug screening study^20^ previously published by Bihler et al. against the FDA-approved list^3^. We considered trafficking efficiencies to focus on misfolding/mis-trafficking variants. This revealed an expanded collection of unapproved and poorly responsive variants also localized in NBD1 and ICL4 (**Supplemental Figure S1D**). Additional poorly responsive variants occur in the lasso motif, the N-terminus of TMD1, the middle of TMD2, and NBD2. Finally, since F508del folds at lowered temperature^21^, we filtered a previous temperature sensitivity screen from Anglés et al. against the FDA-approved list^16^ (**Supplemental Figure S1E**). Poor responders V520F, R560T, R560K, and L1065P were rescued by temperature correction, while L558S, A559T, Y569D, and R1066C were not. Differential temperature correction profiles indicated poorly responsive variants diverge in rescue susceptibility.

To confirm response profiles, We tested selectively and poorly responsive variants in a HEK293T transient expression system. Briefly, CFTR expression plasmids encoding WT and variants were transfected into HEK293T cells and treated for 24 h with 3 μM VX-661, 3 μM VX-445, 3 μM VX-445 + 3 μM VX-661 combination, or DMSO vehicle. Lysates were resolved via SDS-PAGE and immunoblotted for CFTR. Core-glycosylated, ER-resident B band CFTR at 175 kDA and fully glycosylated, post-Golgi C band CFTR at >200 kDA were quantified and used to calculate the trafficking efficiency inferred by the ratio of C band to total CFTR e.g., C/(C+B) ratio. All VX-445 selective variants demonstrated robust CFTR rescue by VX-445 and a moderate rescue by VX-661 (**Figure 1E, Supplemental Figure S1F**). Comparing VX-445 versus VX-661 trafficking efficiency confirmed VX-445 selective response for all variants but Y1032C, which was promiscuously responsive (**Figure 1E**).

Poorly responsive variants L558S, A559T, R560T, A561E, and Y569D showed no trafficking efficiency increase with dual drug treatment (**Figure 1F, Supplemental Figure S1G**). V520F and R1066C, by contrast, showed a slight increase, albeit with <50% WT levels and <F508del response (**Figure 1F**). These data suggest that the response profiles of our variants of interest are robust and reproducible in our transient expression system, amenable to multiplexed quantitative interactomics to characterize proteostasis interactions.

### **II.** Interactome remodeling of responsive and poorly responsive CFTR variants diverge

We integrated our previously established affinity purification mass spectrometry (AP-MS) approach with 18-plex tandem mass tags (TMT) for increased multiplexing to quantify relative interactor levels across datasets – selectively responsive variants with VX-445 correction and poorly responsive variants with VX-661 and VX-445 dual drug correction.

Briefly, HEK293T cells were transiently transfected with CFTR expression plasmids and treated with correctors or vehicle. Lysates were immunoprecipitated for CFTR and interactors with anti-CFTR monoclonal antibody-conjugated Sepharose beads. Proteins were eluted, digested, TMT labeled, and samples pooled for LC-MS/MS analysis (**Supplemental Figure S2A**). Quantitative comparisons were performed after the median normalization of TMT intensity to remove non-specific interactors (**Supplemental Figure S3A-B, see Methods**). Interactors were identified by statistically comparing CFTR affinity purification protein levels to mock transfection control protein levels to remove non-specific proteins bound to the beads.

First, we determined the interactomes of five VX-445 selectively responsive variants (**Supplemental Figure S2B**). We included L165S (n=5), Y1032C (n=6), T1036N (n=6), H1054D (n=6), R1066H (n=5) in our AP-MS analysis all with ± VX-445 conditions (**Supplemental Dataset S1**). A mock transfection (n=7), WT CFTR (n=7), and F508del ± VX-445 (n=5) were included as controls. CFTR pull-down enriched bait protein and previously well-characterized interactors^22,23^ (**Supplemental Figure S3C-I**). VX-445 elevated CFTR levels for each variant (**Supplemental Figure S4**). We compiled statistically significant interactors for each selectively responsive variant into a master list comprising 655 interactors.

Next, an identical procedure was performed for seven poorly responsive variants (**Supplemental Figure 2C**). We treated poorly responsive variants with the combination of VX-661 and VX-445 (3µM each) to investigate whether poorly responsive variants exhibit any degree of proteostasis remodeling. We included V520F (n=5), L558S (n=6), A559T (n=5), R560T (n=6), A561E (n=6), Y569D (n=5), and R1066C (n=6), and ± VX-445/VX-661 conditions for all variants (**Supplemental Dataset S1**). A mock transfection (n=6), WT CFTR (n=6), and F508del ± VX-44/661 (n=6) were included as controls.

Again, we median-normalized TMT intensities before quantitative comparison to remove non-specific interactors (**Supplemental Figure S5A-B, see Methods**). CFTR samples showed enriched bait protein and previously identified interactors (**Supplemental Figure S5C-K**). Treatment with VX-445/VX-661 failed to change interaction intensities compared to the vehicle (**Supplemental Figure S6**). Again, we compiled statistically significant interactors into a master list comprising 284 interactors.

Overall, the poorly responsive variants revealed a relatively small number of interactors compared to the VX-445 selectively responsive variants. However, we observed an overlap of 105 shared core interactors between both datasets (**Supplemental Figure 2D**). These data suggest selectively and poorly responsive variants share core interactions but experience unique perturbations during correction. We leveraged these interactomics datasets to understand the proteostasis landscapes of CFTR variants during correction.

### **III.** VX-445 attenuates proteostasis interactions across pathways for selectively responsive variants

To understand the proteostasis interactome commonalities and differences amongst the VX-445 selectively responsive variants, interactors were organized into biological pathways annotated based on Gene Ontology (GO) terms (**Supplemental Dataset S3**) and filtered down to 170 interactors involved in protein homeostasis pathways such as translation, protein folding, proteasomal and autophagy degradation, trafficking, and endocytosis. A K-means cluster-map of Spearman correlation coefficients between variants and conditions split variants and conditions into four clusters (**Figure 2A, Supplemental Figure S7A, Supplemental Dataset S2**). These four clusters grouped 1) WT with T1036N 2) L165S, 3) F508del with R0166H, and 4) Y1032C with H1054D (**Figure 2A**). Clustering suggests VX-445 restores T1036N interactome the most towards WT, whereas R1066H likely experiences similar interactions to F508del.

**Figure 2.**
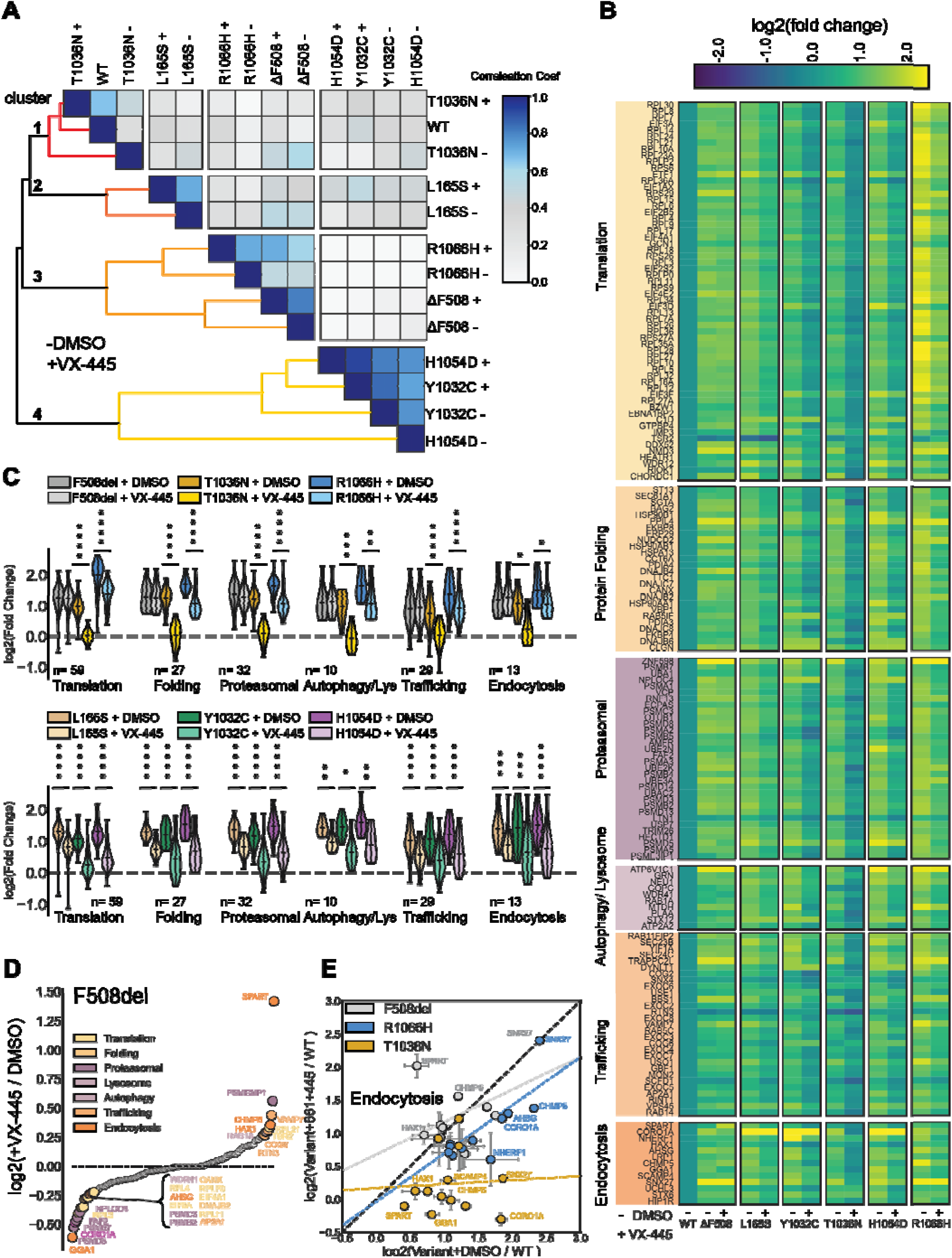
Proteostasis landscapes of VX-445 selectively responsive variants reveal interaction attenuation across pathways. **A.** Ranked Spearman correlation coefficient cluster-map of 170 proteostasis interactions correlations between VX-445 selectively responsive variants. Ordered into four clusters by K means clustering. **B.** Heatmap depicting interactions log2 fold changes of variants normalized to WT CFTR and organized by pathways from GO-term analysis. Yellow represents interactions that are upregulated compared to WT, blue represents interactions that are downregulated compared to WT. **C.** Violin plots of protein enrichment distribution levels for proteostasis pathways from **B.** with known consequences for CFTR. Side-by-side comparisons under DMSO (vehicle) and VX-445 (3 μM) treatment conditions are shown. Statistical significance was calculated using a one-way ANOVA with Geisser-Greenhouse correction and Tukey post-hoc multi-hypothesis correction, and adjusted p values were depicted by *< 0.05, **< 0.01, ***< 0.001, and ****< 0.0001. **D.** Waterfall plot of the log2 fold change between F508del + 3 μM VX-445 over F508del + DMSO organized by rank order of fold change. Interactors above zero are stronger under drug treatment, and interactors under zero are greater in the basal state. Interactors deviating more than one-half standard deviation from the distribution are colored by pathway. **E.** Correlation of individual endocytosis protein quantification between DMSO and VX-445 treatment for F508del (grey), T1036N (gold), and R1066H (blue). A normal line (black dotted line) represents x = y, i.e. no change with the drug. A least squared linear best fit dotted line colored for each variant represented the off-normal axis behavior during drug treatment.

We compared the log2 fold change of individual protein interactors using the WT as a reference. VX-445 decreased interactions across all variants, with Y1032C and T1036N displaying the greatest shifts (**Figure 2B, Supplemental Dataset S4**). Selectively responsive variants interacted with a diverse suite of ribosomal subunits, proteasomal subunits, co-chaperones such as DNAJB4/6, trafficking factors such as COG2/6 and EXOX2/3/4/6/8, and endocytosis machinery such as HAX1. VX-445 decreased interactions with previously characterized CFTR folding and degradation factors such as CANX^22–24^, BAG2^25,26^, DNAJB2^27^, FKBP4/8^28^, PSMC3^13^, Hsp70^29,30^, Hsp90^12,31,32^ isoforms, and NHERF1^33^ (**Figure 2B**). Basal R1066H strongly interacts with translation machinery, suggesting the R1066H variant is evaluated co-translationally (**Figure 2B**).

The distribution of interaction changes for individual pathways provides insights into how VX-445 modulates proteostasis pathways in aggregate. For clarity, we split the variants into a F508del, T1036N, and R1066H plot and a L165S, Y1032C, and H1054D plot depicting both the basal and VX-445 treated conditions (**Figure 2C, Supplemental Dataset S5**). The CFTR-relevant proteostasis pathways were arranged in approximate order of biogenesis: translation, folding, proteasomal degradation, lysosomal degradation/ autophagy, trafficking, and endocytosis - although we note these processes occur simultaneously in cells. VX-445 significantly decreased interactions across all proteostasis pathways for selectively responsive variants (**Figure 2C**).

Surprisingly, VX-445 failed to alter F508del interactions (**Figure 2C**). VX-445 increased F508del interactions with late biogenesis proteins (endocytosis factors SPART, CHMP5, and HAX1 and trafficking factors VAMP7, COG6, and RTN3) while decreasing interactions with endocytosis factors GGA1 and CORO1A (**Figure 2D**). Individual endocytosis interactor correlations between DMSO and VX-445 were examined (**Figure 2E**). The best-fit lines (colored dotted) imply off-axis divergence from basal interaction levels. VX-445 diverted F508del and T1036N endocytosis factors (**Figure 2E**). Indeed, endocytosis represents the most diverted F508del pathway among proteostasis pathways (**Supplemental Figure S7B-F**). These data suggest that VX-445 may modulate F508del interactions with later-stage biogenesis.

### **IV.** Correctors guide poorly responsive CFTR variants toward productive proteostasis landscapes

Again, we filtered poorly responsive interactors for 65 protein homeostasis factors (**Supplemental Dataset S3**). A K-means cluster map of pairwise Spearman correlation coefficients split variants and conditions into 5 clusters (**Figure 3A, Supplemental Figure S8A, Supplemental Dataset S2**,). These clusters grouped F508del + VX-445/661 with WT (cluster 4), Y569D, R1066C, and A561E, regardless of treatment (cluster 5) R560T + DMSO with L558S and A559T + VX-445/661 (cluster 1), L558S + DMSO with R560T + VX-445/661 and (cluster 2) F508del, V520F, and A559T + DMSO with V520F + VX-445/661 (cluster 3).

**Figure 3.**
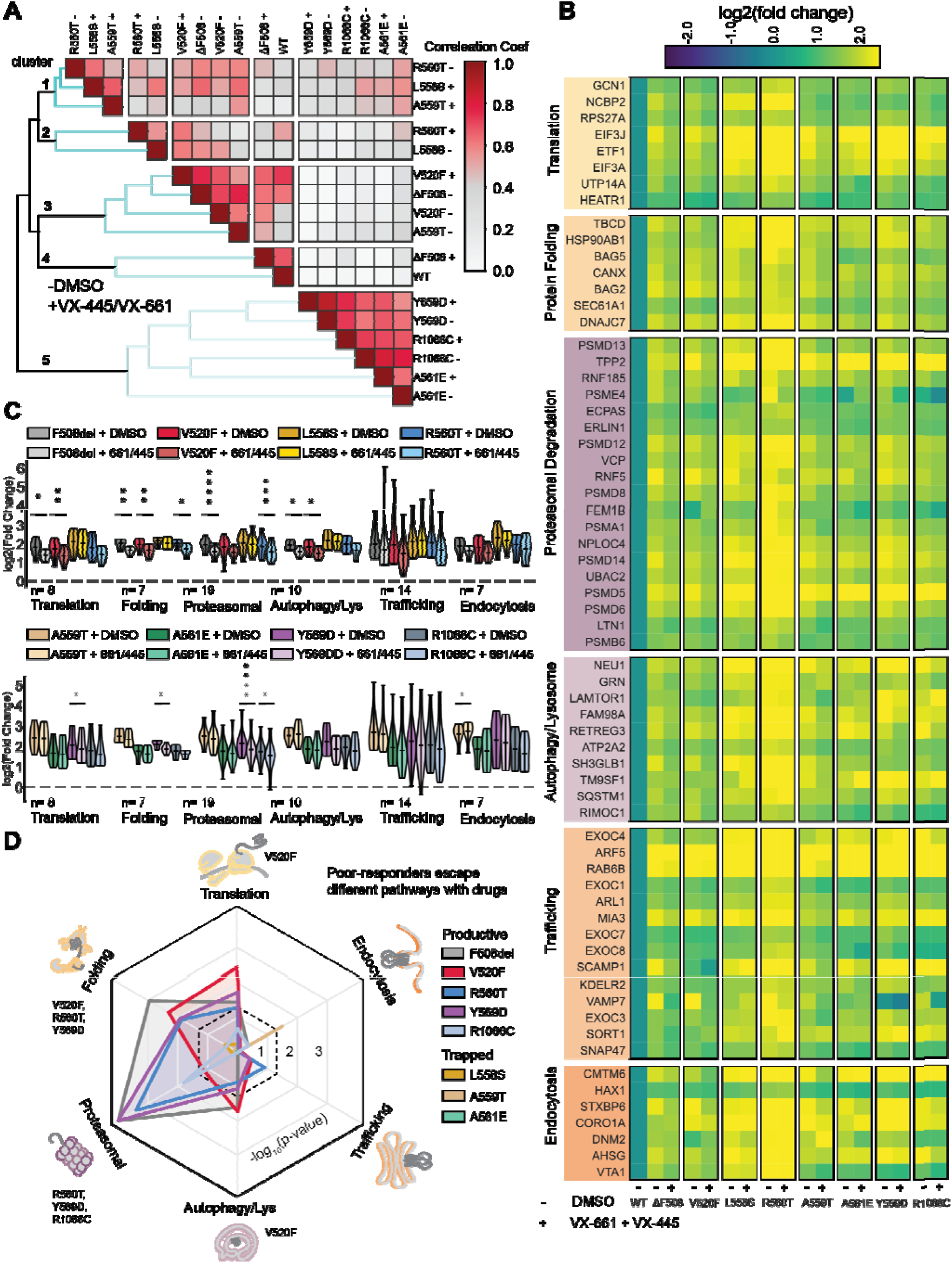
Correctors subdivide poorly responsive variants into productive or trapped proteostasis states. **A.** Spearman ranked correlation coefficient cluster-map for 65 proteostasis interactions correlated between poorly responsive variants. Ordered into five clusters using K means clustering. **B.** Heatmap depicting interactions log2 fold changes of variants normalized to WT CFTR and organized by pathways from GO-term analysis. Yellow represents interactions that are upregulated compared to WT; blue represents interactions that are downregulated. **C.** Violin plots of protein enrichment distribution levels for proteostasis pathways from **B.** under basal and VX-445/VX-661 (3 μM) treatment. Statistical significance was calculated using a one-way ANOVA with Geisser-Greenhouse correction and Tukey posthoc multi-hypothesis correction, and adjusted p values were depicted by *< 0.05, **< 0.01, ***< 0.001, and ****< 0.0001. **D.** Spider plots depicting the -log10 (p values) from the violin plots in **C.** across proteostasis pathways. The black dotted line represents a p-value of 0.05 and serves as a minimum cut-off for statistical significance.

The log2 fold changes of each variant’s interactors, with reference to WT, were compared between vehicle and drug treatment and organized by proteostasis pathways (**Figure 3B, Supplemental Dataset S3**). Several previously known interactors were identified, such as CANX^22–24^, BAG2^25,26^, BAG5^34^, Hsp90AB1^31^, RNF5^35^, and RNF185^36^. Yet, minimal proteostasis changes occurred during dual corrector treatment.

To determine specific proteostasis alterations, we again plotted the aggregate distribution for each pathway in approximate order of biogenesis: translation, folding, proteasomal degradation, lysosomal degradation/ autophagy, trafficking, and endocytosis (**Figure 3C, Supplemental Dataset S6**). VX-661 and VX-445 decreased F508del interactions with translation, protein folding, proteasomal degradation, and lysosomal degradation/ autophagy (**Figure 3C**).

Dual corrector treatment attenuated V520F interactions with translation, protein folding, and lysosome/autophagy machinery. On the other hand, R560T displayed decreased interactions with protein folding and proteasomal degradation factors only (**Figure 3C**). Y569D interactions with translation and protein folding decreased, while both Y569D and R1066C interactions with proteasomal degradation decreased (**Figure 3C**). Dual corrector treatment failed to change interaction levels for L558S with any pathway, consistent with its kinetically trapped misfolding^37^. Correctors also failed to alter A559T and A561E proteostasis interactions. These results indicate that V520F, R560T, Y569D, and R1066C move towards *productive* proteostasis while L558S, A559T, and A561E remain *trapped* during correction.

Even trapped variants show subtle differences. Drugs decreased L558S and A561E interactions with autophagy protein SH3GLB1 – which may recruit proteins to membranes with high curvature and promote membrane fusion^38^ - but increased A559T interactions with SH3GLB1 (**Supplemental Figure S8B-E**). This suggests dual correction pushes L558S and A561E to a proteostasis checkpoint that A559T reaches basally.

The statistical significance of proteostasis pathway remodeling splits the poorly responsive variants into *productive* and *trapped* proteostasis variants (**Figure 3D**). R560T and Y569D demonstrate a distinct pattern of interactions with autophagy/lysosomal and translation proteins. L558S, A559T, and A561E demonstrated no statistically significant proteostasis landscape remodeling – again suggesting trapped proteostasis (**Figure 3D**). Thus, some poorly responsive variants escape proteostasis evaluation during correction and are guided toward productive proteostasis, while others remain trapped despite correctors.

### **V.** VX-445 fails to correct NBD1 structural defects in trapped proteostasis variants

The relationship between proteostasis landscapes and structural defects amongst CFTR variants remains unclear. Experimentally determined structures are only available for F508del; hence, the structural defects of other variants are only accessible through computational modeling. Using our previously established^39^ and applied^10^ Rosetta Comparative Modeling (RosettaCM) method, we determined the structural defects of selectively and poorly responsive CFTR variants.

Briefly, we refined an ensemble of CFTR models into active (PDB 6MSM)^40^ and inactive (PDB 5UAK)^41^ CFTR cryo-EM density maps using automated refinement in Rosetta^42^. Using two conformations distinguishes how variants affect one state over the other and increases conformational sampling during simulation. We then selected the lowest Rosetta scoring models and docked VX-445 to a binding pocket determined by Cryo-EM^8^. Prior focus on Type-I correctors^15,39^ inspired us to pursue variants bound to Type-III corrector VX-445. We used the same coordinates for docked and apo models, so energetic calculations were directly comparable. *In silico* mutagenesis on the apo and VX-445 docked models were performed for both the active and inactive states. We then generated 1000 structural models with RosettaCM for each variant.

The Rosetta score of the 100 lowest-scoring apo models (10% of the total generated models) was plotted against the lowest-scoring 100 VX-445 models. VX-445 globally stabilized poorly responsive variants in the active conformation (**Supplemental Figure S9A**). By contrast, in the inactive conformation, VX-445 only stabilized V520F, Y569D, and R1066C but not L558S, A559T, R560T, and A561E (**Supplemental Figure S9B**).

To understand variant-specific structural defects, we mapped the ensemble change in root-mean-square deviation (ΔRMSD) between WT and Y569D apo state onto the inactive structure (5UAK) (**Figure 4A**). This represents the variant structural instability. We compared it to the ΔRMSD between the apo and VX-445 bound state, representing VX-445 conferred stability (**Figure 4A**). A large positive ΔRMSD (red) suggests destabilizing effects of mutation, whereas a large negative ΔRMSD (blue) suggests stabilizing effects of VX-445. The ΔRMSD values were summed across the region and rolled in a bar plot to compare the variant structural instability versus VX-445 conferred stability (**Figure 4B**).

**Figure 4.**
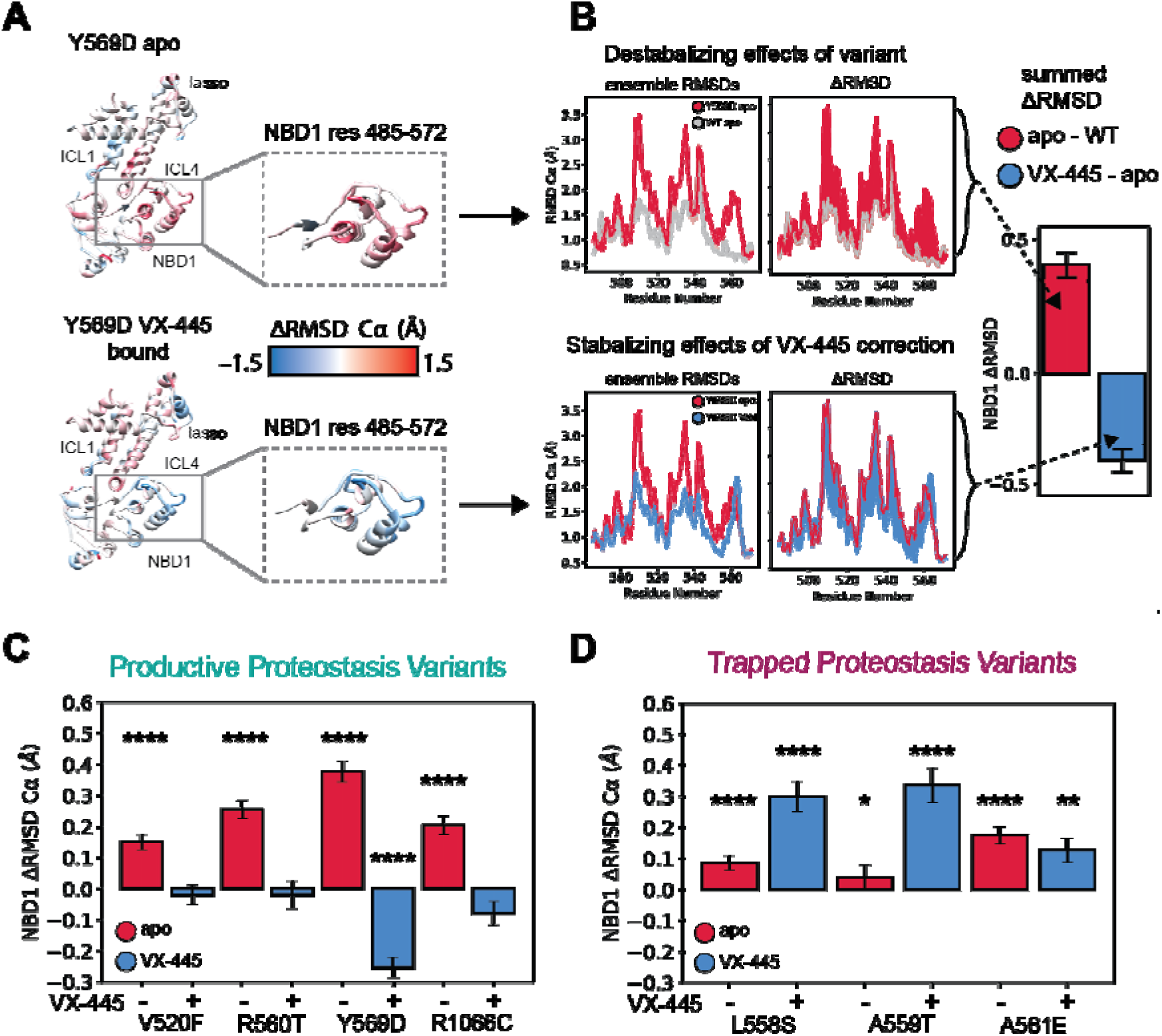
VX-445 fails to stabilize NBD1 of trapped proteostasis poorly responsive variants. **A.** Comparison of variant instability and VX-445 conferred stability. Top: the ΔRMSD between apo Y569D and WT CFTR in the inactive conformation was mapped onto PDB ID 5UAK^41^. Bottom: the ΔRMSD between VX-445 bound Y569D and apo Y569D CFTR in the inactive conformation was mapped onto PDB ID 5UAK^41^. Intercellular Loop 1 (ICL1), ICL4, Lasso motif, and NBD1 are shown for clarity. **B.** Quantification of average ΔRMSD in NBD1 residues 485-572 as an example calculation. Top: the ensemble RMSD by residue is shown for apo Y569D and WT CFTR, followed by a plot showing the ΔRMSD between the RMSDs shaded in red. This represents the instability caused by the Y569D mutation in NBD1. Bot om: the ensemble RMSD by residue is shown for VX-445 bound Y569D and apo Y569D CFTR, followed by a plot showing the ΔRMSD between the RMSDs shaded in blue. This represents the stability conferred by VX-445 to Y569D NBD1. ΔRMSD values are summed in the quantification bar plot portrayed below. **C.** Summed ΔRMSD of inactive conformation NBD1 for poorly responsive variants in productive proteostasis states under drug treatment. Error bars represent the standard error of the mean. Colored as in **B.** **D.** Summed ΔRMSD of inactive conformation NBD1 for poorly responsive variants in a trapped proteostasis under drug treatment. Error bars represent the standard error of the mean. Colored as in **B.** Statistical significances were calculated using a non-parametric Wilcoxon signed-rank test compared to zero to determine if distributions were significantly different from zero, and p values were depicted by *< 0.05, **< 0.01, ***< 0.001, and ****< 0.0001.

NBD1 structural defects differed amongst the poorly responsive variants in productive vs. trapped proteostasis states (**Figure 4C-D, Supplemental Figure S9C-I**). VX-445 reduced the NBD1 ΔRMSD for productive proteostasis variants V520F, R560T, and Y569D (**Figure 4C**), while either not reducing or increasing NBD1 ΔRMSD for the trapped proteostasis variants (**Figure 4D**). This suggests NBD1 remains unstable during VX-445 treatment for trapped proteostasis variants.

We performed the same structural modeling and analysis for the VX-445-selective responders (**Supplemental Figure S10-S11**). VX-445 broadly stabilized the active conformation while failing to stabilize the inactive conformation (**Supplemental Figure S10A-B, Supplemental Dataset S7**), suggesting NBD dimerization may facilitate correction. All selective responders destabilize ICL4 in the active conformation with VX-445, decreasing the ΔRMSD in ICL4 more than in NBD1 (**Supplemental Figures S10C-F, S11A-E**). The inactive conformation shows similar shifts in ICL4 ΔRMSD compared to NBD1 ΔRMSD (**Supplemental Figures S10G-H, S11F-J**). These data suggest the energetic contributions of VX-445 to responders dominate in the active conformation and that VX-445 likely corrects by stabilizing ICL4 over NBD1.

### **VI.** Stabilizing NBD1 rescues poorly responsive variant trafficking

The structural modeling data posited that trapped poorly responsive variants lack NBD1 stability despite VX-445 binding. Thus, we hypothesized that NBD1 stabilizing secondary mutations may rescue poorly typically combined multiple secondary mutations to stabilize CFTR, obstructing the assessment of their individual energetic contributions. Thus, we introduced stabilizing mutations individually.

We chose three NBD1 stabilizing mutations, F494N, I539T, and Q637R, with distinct stabilization mechanisms (**Figure 5A**). F494N stabilizes the Q-loop (residues 491-500), a region involved in ATP binding^44^, I539T stabilizes the structurally diverse region (SDR – residues 533-548)^45^, and Q637R allosterically stabilizes the ICL4 binding pocket near F508^46^. We introduced these stabilizing mutations into three poorly responsive variants, A559T, R560T, and Y569D, as well as into F508del (as a control), then measured drug response.

**Figure 5.**
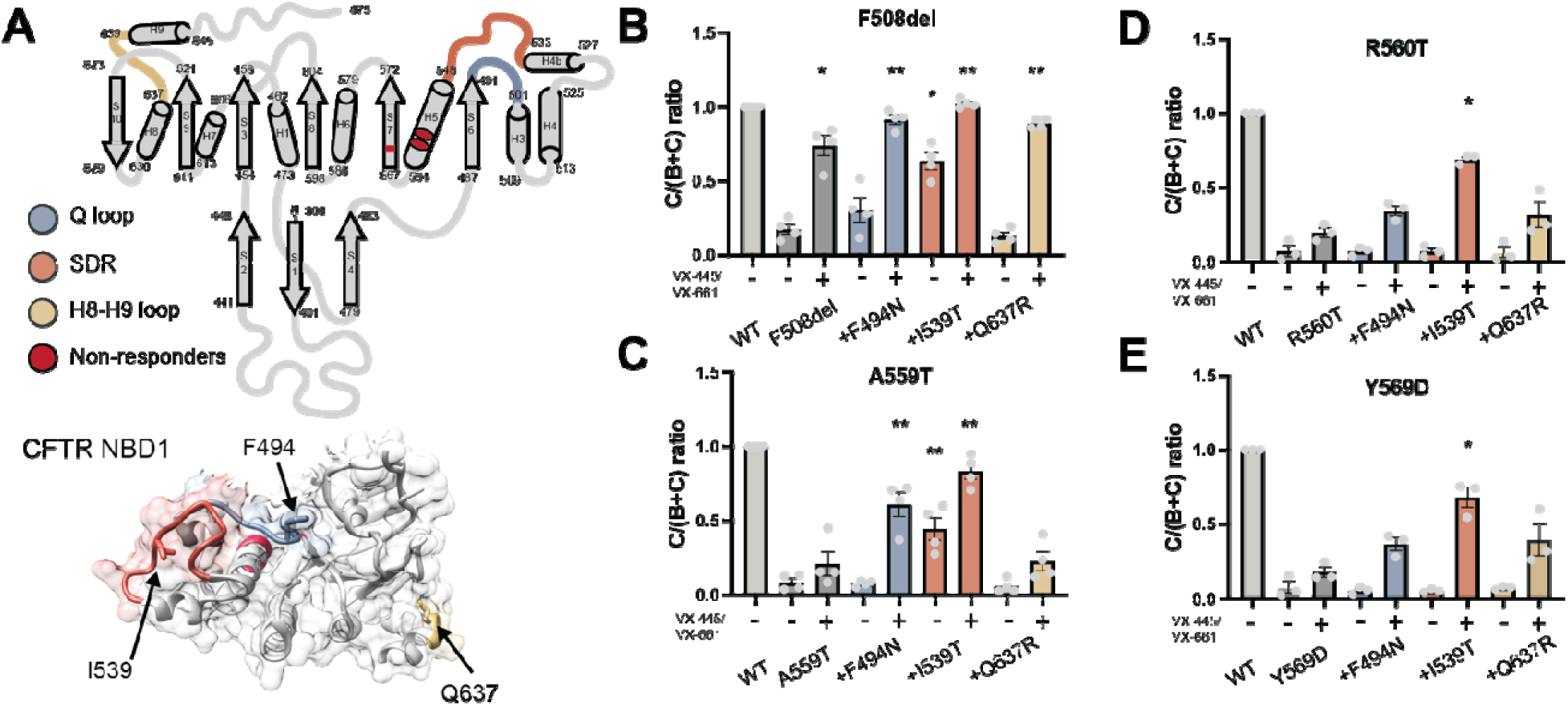
Stabilizing NBD1 with secondary mutations rescues poorly responsive variant trafficking. **A.** Top: ribbon diagram of the CFTR NBD1 structure with poor responders and loops susceptible to stabilizing secondary mutations highlighted. The Q-loop (residues 491-500) shown in blue, the SDR (residues 533-548) shown in orange, and the helix 8 – helix 9 (H8-H9) loop (residue 636-639) shown in gold. Bottom: CFTR NBD1 folded structure viewed from the NBD dimerizing face (5UAK^41^), F494, I539, and Q637 are depicted in their respective loops. The SDR and the Q-loop lie proximate to the poorly responsive variants (red), whereas the H8-H9 loop lies on the other side of NBD1. **B.** Quantification of C/(C+B) band ratio demonstrates dual 3 uM VX-445/VX-661 treatment combined with all secondary mutations F494N, I539T, and Q637R rescue F508del. **C.** Quantification of C/(C+B) band ratio demonstrates secondary mutations F494N or I539T plus dual corrector treatment rescues A559T. **D-E.** Quantifying the C/(C+B) band ratio demonstrates secondary mutations I539T combined with drug treatment rescues R560T and Y569D trafficking. Statistical significances for **B.**-**E.** were calculated using a parametric ratio paired t-test, and p values were depicted by *< 0.05, **< 0.01, ***< 0.001, and ****< 0.0001. Representative blots are shown in **Supplemental Figure S12A-D**.)

Dual drug treatment or I539T alone rescued F508del CFTR, while F494N and Q637R rescued only in combination with correctors (**Figure 5B, Supplemental Figure S12A**). I539T plus dual correctors showed the most robust increase in trafficking efficiency for the three poorly responsive variants (**Figure 5C-E, Supplemental Figure S12B-D**). A559T trafficking was also rescued by combining drug treatment with F494N (**Figure 5C, Supplemental Figure S12B**). These results suggest additional NBD1 stability beyond VX-661/VX-445 is necessary to rescue poor responders.

### **VII.** Distinct proteostasis signatures demarcate CFTR drug vulnerability

To compare proteostasis between selectively and poorly responsive variants, we determined an overlap of 31 proteostasis factors between the datasets (**Supplemental Figure 13A, Supplemental Dataset S8**). Pearson correlation between the datasets of F508del + DMSO, the only conditions found in both, was 0.72, and the Spearman correlation was 0.68 (**Supplemental Figure 13B**). Fair correlation indicated the two datasets show similar F508del interaction levels but not exact correspondence.

Principal component analysis (PCA) of the 31 proteostasis factors revealed that the first principal component accounts for 83% of the variance amongst the interactors (**Figure 6A**). Responsive variants shift substantially in the first principal component (ΔPC1) (**Figure 6B**), suggesting PC1 captures proteostasis changes driving response. By contrast, poorly responsive variants show little shift in PC1 but a shift in PC2 (**Supplemental Figure 13C**). Notably, opposing axes shifts played out in t-SNE and UMAP dimensionality reduction (**Supplemental Figure 13D-E**), demonstrating this not an artifact of the dimensionality reduction technique. Specifically, the trapped variants (A559T, A561E, and L558S) ΔPC1 are lower than the productive proteostasis variants (V520F, R560T, Y569D, and R1066C) (**Figure 6B**). The PC1 loadings for all genes were similar, indicating no individual proteostasis factor was driving the PCA (**Figure 6C**). Thus, PC1 captures meaningful information about CFTR drug vulnerability.

**Figure 6.**
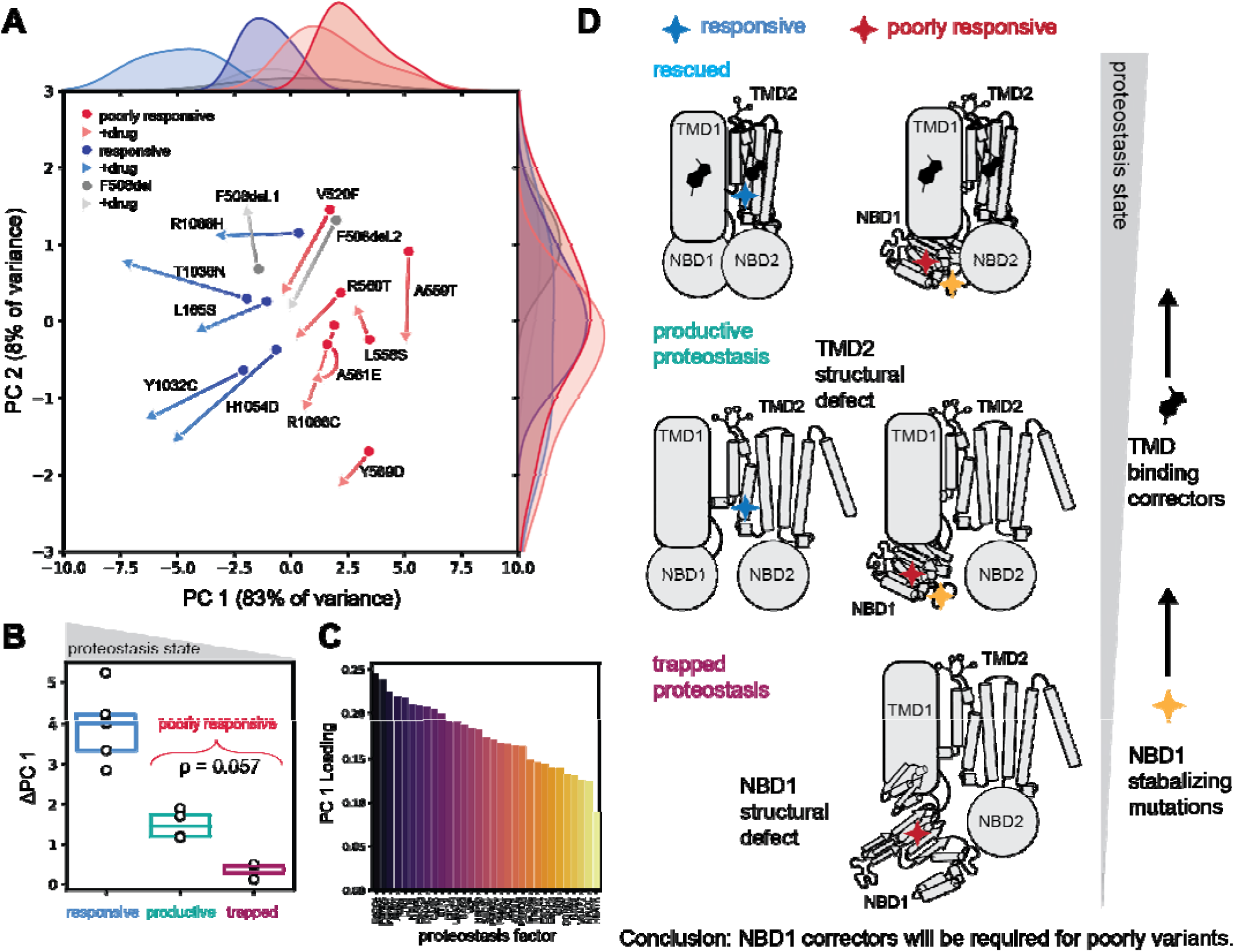
Proteostasis landscapes and structural defects drive CFTR drug response vulnerability. **A.** Principal component analysis of 31 overlapping proteostasis factors between selectively and poorly responsive data sets. Basal conditions are shown as circles, and drug conditions (VX-445 or VX-661+VX-445, respectively) are shown as triangles. A line connecting basal and drug points for the same variants shows how correction shifts interactions for each variant across PC space. **B.** Distributions of the first principal component shift (ΔPC1). **C.** The ordered PC1 loadings demonstrate which proteostasis factors contributed the most to the PCA. **D.** A schematic showing the putative relationship between CFTR proteostasis and structural defects. Selectively responsive variants experience TMD2 structural defects and relatively productive proteostasis; thus, they respond readily to current corrector compounds that bind to the TM domains. Poorly responsive variants, by contrast, experience NBD1 structural defects and relatively unproductive or trapped proteostasis states. By stabilizing NBD1 with secondary mutations, poorly responsive variants can be moved further up the gradient of productive proteostasis and thermodynamic stability to where current TMD binding correctors can rescue them.

NBD1 variants with low ΔPC1 shift also show NBD1 structural defects (**Figure 4C-D**). We propose a model where TMD structural defects observed in selectively responsive variants allow robust drug response. In this model, NBD1 structural defects must be thermodynamically stabilized to move trapped poorly responsive variants toward productive proteostasis and promote rescue with current therapeutics (**Figure 6D**).

## DISCUSSION

Despite the advent of revolutionary corrector compounds for treating CF, 10% of persons with CF have no etiological treatment options, with ∼3% of the total CF population harboring poorly responsive class II variants^4^. Correctors rescue the derailed protein homeostasis and structural stability of CFTR through diverse mechanisms that split response profiles into selectively and poorly responsive variants. Poorly responsive variants remain understudied due their prevalence in underrepresented populations and subsequent low measured allele frequencies. Increased genetic sequencing among underrepresented persons may reveal emerging poorly responsive variants – creating the need to predict response^47^. Here, we sought to understand the proteostatic and structural deficiencies of selectively and poorly responsive variants to reveal opportunities for correction.

Poorly responsive variants occur in under-sampled patient demographics. A559T is common amongst individuals of African descent^48^ and is the second most common mutation (after F508del) in the Dominican Republic^49^. R1066C is the second most common variant (after F508del) in Puerto Rican patients^49^. A few previous *in vitro* studies characterized poorly responsive variants with correctors in various CF model systems. In stably expressing CF bronchial epithelial cells (CFBEs), variants L558S, A559E/T, and R560T failed to show any response to VX-661/445 combination treatment^17^. A559T only showed 10% WT function after treatment with VX-661/VX-445 in homozygous-derived rectal organoids^18^. Variants V520F, A559T, and Y569D failed to respond to temperature correction in HEK293T cells^16^. By contrast, in CFBEs, V520F and Y569D responded modestly to VX-661/VX-445 together but not VX-661 alone, whereas R560T failed to respond regardless of treatment combination^9^. Stabilizing secondary mutations have also been leveraged to rescue poor responders; for example, G550E rescues A561E but not R560T^19^.

The basal state proteostasis landscapes diverge for selective VX-445 responsive variants. Previously, we postulated that common proteostasis states in the basal condition drove corrector response, e.g. the proteostasis network of selective responders was amenable to correction.^13^ Our new results shift this paradigm among responsive variants from common to more diverse basal proteostasis states. However, these diverse basal proteostasis landscapes are convergently corrected across pathways. VX-445 decreased interactions for all pathways amongst responsive variants. For example, R1066H experienced unique interactions with translation proteins that VX-445 reversed. By contrast, VX-445 primarily attenuated late-stage biogenesis factors for F508del and T1036N. Thus, like VX-809^50^, VX-445 can rescue CFTR variants at multiple points in biogenesis in a variant-dependent manner.

Correctors failed to attenuate interactions across pathways in poorly responsive variants. However, we revealed proteostasis landscape remodeling for a subset of pathways. V520F, R560T, Y569D, and R1066C CFTR exhibit remodeling of early-stage biogenesis such as translation, folding, and degradation pathways when treated with VX-661 and VX-445. Interestingly, previous studies showed V520F and R560T, like F508del, are correctable by temperature reduction (**Supplement Figure S1E**). Despite their similarities, corrector treatment attenuates V520F interactions with autophagy versus attenuating R560T and Y569D interactions with proteasomal degradation, suggesting R560T and Y569D structural defects occur early while CFTR is still amenable to retro-translocation. It remains unclear whether poorly responsive variants, such as R560T, aggregate intracellularly before autophagy degradation, however, this phenomenon was observed for N1303K CFTR^51^.

VX-661 failed to stabilize A559T, R560T, and L558S in previous FoldX simulations^17^. Here, VX-445 failed to stabilize L558S, A559T, and A561E. We hypothesized that enhanced NBD1 stabilization could rescue poorly responsive variants. In combination with dual correction treatment, SDR stabilizing I539T consistently rescued poor responders. However, it remains unclear if I539T energetic contributions (on the order of -2.5°C ΔT_m_ ^39,52^) or location in the SDR is more important for rescuing NBD1 core variants. NBD1 correctors already in development^53^ or next-generation correctors^54^ may be used to treat NBD1 poor responders characterized here.

Combining orthogonal proteostasis profiling with structural simulations reveals distinct albeit related information about CFTR corrector response. Interactomics divided poor responders into productive proteostasis showed NBD1 structural defects in Rosetta simulations with VX-445 bound. Despite the observed correspondence for proteostasis and structural defect, strong thermodynamic stabilizing secondary mutations rescue all poorly responsive variants tested. We propose a model in which poorly responsive NBD1 variants are trapped in unproductive proteostasis states that can only be alleviated by introducing thermodynamic stability directly to NBD1 (**Figure 6D**).

We note some important limitations. Proteostasis factors can vary in expression between cell lines. However, the proteostasis network is conserved at the pathway level e.g. proteasomal proteins and autophagy proteins between HEK293T cells and other CF-derived model systems. Thus, we assume the pathway-level analysis will hold across model systems. Furthermore, CFTR response correlates between HEK293T cells and CFBEs^16^, and we observed consistent response profiles in our system (**Figure 1E-F**). Interactors identified in HEK293T cells and then knocked down in CFBE cells sensitized P67L to VX-445^14^, highlighting the functional conservation of proteostasis factors across cell lines. Interactome changes do not necessarily map one-to-one with CFTR correction due to redundancies in proteostasis components. In our PCA, PC1 loadings were similar across proteostasis factors (**Figure 6C**), suggesting no single interaction drives drug response. Our computational modeling only samples conformational space far from the folding equilibrium process. We note some variants may never reach such folded conformations. However, correction suggests misfolding-prone CFTR variants sample near folded conformations in cells during rescue, although they may comprise a small fraction of the conformational ensemble.

Undersampling of persons with CF worldwide limits our understanding of the true allelic frequency of poorly responsive variants. Changing patient demographics and the decreasing price of genetic sequencing may change the distributions of alleles in the CF population. This will require efficient assessment strategies for variant responses to current and future correctors to expand treatment to all persons with CF. We show that proteostasis profiling and computational structural modeling provide a framework for discerning elusive drug response mechanisms. Subtle differences in proteostasis landscape changes among poor responders were accompanied by underlying structural defects among the same variants. This approach may be leveraged by future investigations seeking to expand therapeutics to new or unapproved variants associated with other protein misfolding diseases in broad personalized medicine applications.

## METHODS

### Bioinformatic analysis

The McKee et al., Deep Mutational Scan (DMS) dataset was downloaded from^10^– specifically Supplementary Table S1. The cell surface immunostaining intensity was normalized to WT (1695 units). The basal state DMSO vehicle and drug response VX-445 and VX-661 data was split to plot the corrector response. Variants demonstrating >100% WT immunostaining intensity were filtered out to consider poorly responsive variants. The Bihler et al. dataset, Supplementary Table S1, was downloaded from^20^. Only trafficking data of missense variants in the DMSO and VX-445/661 conditions were considered to focus on variants amenable to correctors. Variants with <25% WT C/(C+B) band ratio under dual drug treatment conditions were considered poorly responsive. The Anglés et al. dataset was downloaded from^16^. Again, trafficking data only was used. The standard temperature trafficking index (TrIdx 37) and the reduced temperature corrected trafficking index (TrIdx 27) were compared. Variants with >90% WT temperature correction were removed. The variants from each study were filtered against an FDA-approved variant list from Vertex Pharmaceuticals^3^ (https://www.trikafta.com/who-trikafta-is-for, accessed on 08/2023) and the response was plotted as a function of CFTR sequence.

### Plasmids and antibodies

CFTR expression plasmids in the pcDNA5 vector were gifted from E. Sorscher and J. Hong (Emory University, Atlanta, Georgia)^55^. Site-directed mutagenesis with Q5 polymerase (New England BioLabs) was used to introduce VX-445 selective and poorly responsive mutations in WT CFTR expression vectors. KLD enzyme reaction (New England BioLabs) ligated the PCR product. All plasmids were confirmed by Sanger sequencing of the local region (Azenta).

CFTR immune blotting was performed with anti-CFTR 217 R-domain mouse monoclonal antibodies (http://cftrantibodies.web.unc.edu/) that were purchased from J. Riordan (University of North Carolina, Chapel Hill, North Carolina – The Cystic Fibrosis Molecular/Functional Measurement Core). CFTR co-immunoprecipitation was performed with 24-1 anti–C-terminus monoclonal antibodies (ATCC HB-11947) as previously described^56^.

Antibodies were used in immunoblotting buffer (5% bovine serum albumin [BSA] in Tris-buffered saline, pH 7.5, 0.1% Tween-20, and 0.1% NaN_3_) at 1:1000 dilutions of each primary antibody. Fluorescent secondary antibodies were used at the indicated dilutions in 5% milk in Tris-buffered saline (TBS), 0.1% Tween-20 (TBS-T): goat anti-mouse Starbright700 (1:10,000, Bio-Rad), anti-rabbit rhodamine-conjugated tubulin (1:10,000, Bio-Rad). Immunoblotted PVDF membranes were rinsed in TBS-T and incubated with secondary antibodies in 5% milk at room temperature for 30 minutes, rinsed in TBS-T again, and imaged using a ChemiDoc MP Imaging System (BioRad).

### Cell culture, immunoprecipitation, SDS–PAGE, and immunoblotting

HEK293T cells were maintained in Dulbecco’s Modified Eagle’s Medium of high glucose with 10% fetal bovine serum, 1% penicillin/streptomycin, and 1% glutamine. Cells were transiently transfected 24 hours after seeding 2 × 10^5^ cells/mL with CFTR expression plasmids of respective variants using the calcium phosphate method^57^. Cells were treated 16-18 hours post-transfection with either DMSO vehicle, 3 μM VX-445 for the selectively responsive variants, or 3 μM VX-445 + 3 μM VX-661 for the poorly responsive variants. 24 hours after treating cells, approximately 10^7^ cells were collected for each condition. Cells were lysed and harvested from the plate by incubating in TN buffer with 0.5% IGEPAL CA-630 and a Mini, EDTA-free Protease Inhibitor Cocktail (Roche) cocktail at 4 °C for 20 minutes. Cells were then collected, sonicated for 3 minutes, and centrifuged at 18,000 g for 15-30 minutes. Total protein concentration per sample was normalized using a Pierce BCA Protein Assay kit (Thermo Scientific).

Co-immunoprecipitation protein identification technology (CoPIT), developed by Pankow et al., was utilized for the direct affinity purification of CFTR and its interacting proteins, as previously outlined^56^. In brief, cell lysates were pre-cleared with 100 μL 4B Sepharose beads (Sigma) at 4°C for 1 hour. Precleared lysates were immunoprecipitated overnight using 100 μL slurry (50% beads/50% buffer) of 24-1 anti-CFTR mAB conjugated Protein G sepharose beads at 4°C. Beads were rinsed thrice with TNI buffer, twice with TN buffer, and frozen at -80 °C for at least 1.5 hours. Beads were then eluted twice while shaking at >700 rpm at 37 °C for 30 minutes - 1 hour using a 0.2 M Glycine (pH 2.3), 0.5% IGEPAL CA-630 buffer. The elution was then prepared for mass spectrometry analysis.

### Multiplexed LC-MS/MS

Sample preparation for multiplexed LC-MS/MS was performed as previously described^13,14^. Digested peptide samples were TMT-labeled with TMT Pro 16-plex reagents or 18-plex reagents (Thermo Fisher) in 40% v/v acetonitrile for one hour at room temperature (**Supplementary Table 1**). LC-MS/MS analysis was performed using an Exploris 480 (Thermo Fisher) mass spectrometer equipped with an UltiMate3000 RSLCnano System (Thermo Fisher). MudPIT experiments were performed with eight 10 μL sequential injections of 0, 10, 20, 40, 60, 80, and 100% buffer C (500 mM ammonium acetate in buffer A), followed by a final injection of 90% buffer C with 10% buffer B (99.9% acetonitrile, 0.1% formic acid v/v). Each injection step was followed by a 6 minute 2% B to 2 minute 2% - 5% B, 92-minute gradient from 5% to 35% B, followed by a 5-minute gradient from 35%-65% B, and 1 minute 65% - 85%, held for 7 minutes to drop to 2% instantly and held for 17 minutes for a total of 130 minutes at a flow rate of 500 nL/min, on a 20 cm fused silica microcapillary column (ID 100 μM) ending with a laser-pulled tip filled with Aqua C18, 3 μm, 100Å resin (Phenomenex).

Electrospray ionization (ESI) was performed directly from the tip of the column, applying a voltage of 2.2 kV and an ion transfer tube temperature of 275°C. Data-dependent acquisition of MS/MS spectra was performed by scanning from 375-1500 m/z with a resolution of 120,000. Peptides with an intensity above 5.0E3 with charge state 2-7 from each full scan were fragmented by high collision dissociation with pure N_2_ gas using normalized collision energy of 36, with a 0.7 m/z isolation window, 150 ms maximum injection time, at a resolution of 45,000 scanned at a defined first mass at 110 m/z and dynamic exclusion set to 45, and a mass tolerance of 10 ppm. MS data is available for VX-445 selective responsive variants (**Supplemental Dataset S9**) and poorly responsive variants (**Supplemental Dataset S10**).

### Interactome characterization

Peptide identification and protein quantification based on TMT label relative abundance were conducted using Proteome Discoverer 2.4 as previously described^13,14^. To identify interactors, we used median normalized TMT intensity to compare experimental conditions to mock transfection control. We assume that most identified proteins are nonspecific interactors present in similar amounts in both the mock and CFTR-bait Co-IP samples. Median normalization decreases TMT intensity variability across different LC-MS/MS runs and allows data aggregation between runs (**Supplemental Figure S3A-B and S5A-B**).

### Statistical analysis

To determine statistically significant CFTR interactors, we used a two-tailed paired t-test with scipy.stats.ttest_rel (https://docs.scipy.org/doc/scipy/) package in Python to calculate the p-value between the log2 TMT intensity of each protein over the corresponding log2 TMT intensity in the mock-transfected control condition. A volcano plot with a hyperbolic curve y > c/(x− x0), where y is the adjusted p-value, x is the log2 fold change, x0 is the standard deviation (0.5σ or 1σ), and c is the curvature (c = 0.4). We considered proteins greater than the 1σ curve high confidence interactors and proteins greater than the 0.5σ curve to be medium confidence interactors.

For the distribution of interactors depicted as violin plots in Figures 3 and 4, a one-way ANOVA with Geisser-Greenhouse correction and Tukey post-hoc multi-hypothesis correction was used to determine statistical significance. For these statistical calculations, adjusted *p-*values were depicted by *< 0.05, **< 0.01, ***< 0.001, and ****< 0.0001. For RosettaCM-based simulation and calculations, a non-parametric Wilcoxon test was used to determine statistical significance from zero. For Western blots, a parametric ratio-paired t-test was used. For these statistical calculations, p-values are depicted by * < 0.05, **< 0.01, *** < 0.001, and **** < 0.0001.

### Dimensionality Reduction Analysis

Overlap between selectively responsive and poorly responsive variants proteostasis interactors, e.g. the overlap in proteins from the heatmaps portrayed in Figure 2B and 3B, was determined to comprise 31 proteins (**Supplemental Dataset S8**). Correlation between the only condition in both datasets, F508del + DMSO, was determined using Python libraries <u>scipy.stats.pearsonr</u> and <u>scipy.stats.spearmanr</u> for Pearson and Spearman correlation coefficients, respectively. Dimensionality reduction was calculated in Python using <u>sklearn.decomposition.PCA</u> library for principal component analysis, <u>sklearn.manifold.TSNE</u> library for T-distributed Stochastic Neighbor Embedding, and <u>umap</u> library for uniform manifold approximation.

### Pathway analysis

We manually annotated pathways by cross-referencing gene names on UniProt and classifying pathways using Gene Ontology (GO) terms related to Biological Pathway as previously described^13^. Pathway assignments in the selectively responsive and poorly responsive datasets were cross-referenced to ensure consistent assignment of putative pathways in both datasets.

### CFTR Rosetta Comparative Modeling

We modeled CFTR mutants with Rosetta Comparative Modeling (CM) as previously desribed^39^. Briefly, published cryo-EM maps (PDB ID 5UAK^41^ and 6MSM^40^) were refined^42^ and the lowest-scoring five models were selected for docking of VX-445. Conformational ensembles of VX-445 were generated with the BCL ConformerGenerator^58^ and exported Rosetta-readable parameter file for ligand docking. To dock VX-445, we aligned BCL parameterized VX-445 to the same coordinates as the published bound models^8^ in Chimera and saved the aligned corrector coordinates. VX-445 was docked 1000 times using full atom docking simulations with RosettaLigand^59^ for 24 cycles, repacking every cycle. The interface energy scores and non-superimposed root mean squared (rms) deviation from aligned published VX-445 bound models (PDB ID 8EIQ)^8^ were reported. We chose the docked pose with the least squared distance from the lowest scoring model and rms from the published model as a final template for *in silico* mutagenesis. We mutated patient variants into the cryo-EM refined template models MutateResidue mover in Rosetta. F508del was modeled as previously and RosettaCM^60^ was performed as previously described^39^.

### RosettaCM Analysis

We assessed CFTR variant thermodynamic stability using the Rosetta score and the alpha carbon (Ca) root mean squared deviation (RMSD). Variant energy change was calculated by subtracting the average Rosetta score of the apo from the WT ensemble, while VX-445 binding energy change was calculated by subtracting the VX-445 bound from the apo ensemble. RMSD was calculated per residue in reference to the lowest-scoring model in the ensemble as previously described^39^.

ΔRMSD between apo variant and WT CFTR was mapped onto the corresponding CFTR structure (e.g., active or inactive). The ΔRMSD for NBD1 (residues 485-572) were quantified because they contact TMD2 via ICL4. Mutation instability was calculated by subtracting variant RMSD from WT RMSD (**Figure 4C**). Likewise, VX-445 stability was calculated by subtracting VX-445 bound RMSD from apo RMSD for each variant (**Figure 4C**). The summed ΔRMSD values were presented as bar plots.

## ACKNOWLEDGEMENTS

This work was supported by R35 GM133552 (NIGMS) and R01 HL167046 (NHBLI). EFM was supported by a predoctoral fellowship F31 HL162483 (NHLBI) and Chemical-Biology Interface training grant T32 GM065086 (NIGMS). We thank Olivia Paige Black for helping with the material preparation and Asaf Elazar for helpful feedback on the figures.

## AUTHOR CONTRIBUTIONS

Conceptualization: E.F.M., L.P.; Funding acquisition: E.F.M., L.P., J.M.; Experimental Design: E.F.M., L.P.; Experimental Data Collection: E.F.M., M.K., J.O.; Computational Design: E.F.M., J.M.; Computational Data Collection: E.F.M.; Data Analysis: E.F.M.; Writing original draft: E.F.M.; Writing review and editing, E.F.M., M.K., J.O., J.M., L.P. All authors have read and agreed to the published version of the manuscript.

## REFERENCES

1. Veit, G. et al. From CFTR biology toward combinatorial pharmacotherapy: expanded classification of cystic fibrosis mutations. Mol. Biol. Cell 27, 424–433 (2016).

2. McDonald, E. F., Meiler, J. & Plate, L. CFTR Folding: From Structure and Proteostasis to Cystic Fibrosis Personalized Medicine. ACS Chem. Biol. acschembio.3c00310 (2023) doi:10.1021/acschembio.3c00310.

3. Trikafta Prescribing Information. (2024).

4. CF Foundation Patient Registry, https://www.cff.org/medical-professionals/patient-registry. (2022).

5. Riordan, J. R. et al. Identification of the Cystic Fibrosis Gene: Cloning and Characterization of Complementary DNA. Science 245, 1066–1073 (1989).

6. Fiedorczuk, K. & Chen, J. Mechanism of CFTR correction by type I folding correctors. Cell 185, 158–168.e11 (2022).

7. Veit, G. et al. Structure-guided combination therapy to potently improve the function of mutant CFTRs. Nat. Med. 24, 1732–1742 (2018).

8. Fiedorczuk, K. & Chen, J. Molecular structures reveal synergistic rescue of Δ508 CFTR by Trikafta modulators. Science 378, 284–290 (2022).

9. Veit, G., et al. Allosteric folding correction of F508del and rare CFTR mutants by elexacaftor-tezacaftor-ivacaftor (Trikafta) combination. JCI Insight 5, e139983 (2020).

10. McKee, A. G. et al. General trends in the effects of VX-661 and VX-445 on the plasma membrane expression of clinical CFTR variants. Cell Chem. Biol. 30, 632–642.e5 (2023).

11. Pankow, S. et al. Δf508 CFTR interactome remodelling promotes rescue of cystic fibrosis. Nature 528, 510– 516 (2015).

12. Wang, X. et al. Hsp90 Cochaperone Aha1 Downregulation Rescues Misfolding of CFTR in Cystic Fibrosis. Cell 127, 803–815 (2006).

13. McDonald, E. F., Sabusap, C. M. P., Kim, M. & Plate, L. Distinct proteostasis states drive pharmacologic chaperone susceptibility for cystic fibrosis transmembrane conductance regulator misfolding mutants. Mol. Biol. Cell 33, ar62 (2022).

14. Kim, M. et al. Elexacaftor/VX-445–mediated CFTR interactome remodeling reveals differential correction driven by mutation-specific translational dynamics. J. Biol. Chem. 299, 105242 (2023).

15. McKee, A. G. et al. General trends in the effects of VX-661 and VX-445 on the plasma membrane expression of clinical CFTR variants. Cell Chem. Biol. 30, 632–642.e5 (2023).

16. Anglès, F., Wang, C. & Balch, W. E. Spatial covariance analysis reveals the residue-by-residue thermodynamic contribution of variation to the CFTR fold. *Commun*. Biol. 5, 356 (2022).

17. Zacarias, S., Batista, M. S. P., Ramalho, S. S., Victor, B. L. & Farinha, C. M. Rescue of Rare CFTR Trafficking Mutants Highlights a Structural Location-Dependent Pattern for Correction. Int. J. Mol. Sci. 24, 3211 (2023).

18. Kleinfelder, K. et al. Theratyping of the Rare CFTR Genotype A559T in Rectal Organoids and Nasal Cells Reveals a Relevant Response to Elexacaftor (VX-445) and Tezacaftor (VX-661) Combination. Int. J. Mol. Sci. 24, 10358 (2023).

19. Roxo-Rosa, M. et al. Revertant mutants G550E and 4RK rescue cystic fibrosis mutants in the first nucleotide-binding domain of CFTR by different mechanisms. Proc. Natl. Acad. Sci. U. S. A. 103, 17891– 17896 (2006).

20. Bihler, H. et al. In vitro modulator responsiveness of 655 CFTR variants found in people with cystic fibrosis. J. Cyst. Fibros. S1569199324000213 (2024) doi:10.1016/j.jcf.2024.02.006.

21. Denning, G. M. et al. Processing of mutant cystic fibrosis transmembrane conductance regulator is temperature-sensitive. Nature 358, 761–764 (1992).

22. Okiyoneda, T. et al. Role of calnexin in the ER quality control and productive folding of CFTR; differential effect of calnexin knockout on wild-type and ΔF508 CFTR. Biochim. Biophys. Acta - Mol. Cell Res. 1783, 1585–1594 (2008).

23. Farinha, C. M. & Amaral, M. D. Most F508del-CFTR Is Targeted to Degradation at an Early Folding

24. Rosser, M. F. N., Grove, D. E., Chen, L. & Cyr, D. M. Assembly and Misassembly of Cystic Fibrosis Transmembrane Conductance Regulator: Folding Defects Caused by Deletion of F508 Occur Before and After the Calnexin-dependent Association of Membrane Spanning Domain (MSD) 1 and MSD2. Mol. Biol. Cell 20, 2673–2683 (2008).

25. Arndt, V., Daniel, C., Nastainczyk, W., Alberti, S. & Höhfeld, J. BAG-2 Acts as an Inhibitor of the Chaperone-associated Ubiquitin Ligase CHIP. Mol. Biol. Cell 16, 5891–5900 (2005).

26. Dai, Q. et al. Regulation of the Cytoplasmic Quality Control Protein Degradation Pathway by BAG2. J. Biol. Chem. 280, 38673–38681 (2005).

27. Westhoff, B., Chapple, J. P., Van Der Spuy, J., Höhfeld, J. & Cheetham, M. E. HSJ1 is a neuronal shuttling factor for the sorting of chaperone clients to the proteasome. Curr. Biol. 15, 1058–1064 (2005).

28. Hutt, D. M. et al. FK506 binding protein 8 peptidylprolyl isomerase activity manages a late stage of cystic fibrosis transmembrane conductance regulator (CFTR) folding and stability. J. Biol. Chem. 287, 21914– 21925 (2012).

29. Chiaw, P. K. et al. Hsp70 and DNAJA2 limit CFTR levels through degradation. PLoS ONE 14, 1–27 (2019).

30. Choo-Kang, L. R. & Zeitlin, P. L. Induction of HSP70 promotes ΔF508 CFTR trafficking. Am. J. Physiol. - Lung Cell. Mol. Physiol. 281, 58–68 (2001).

31. Coppinger, J. A. et al. A chaperone trap contributes to the onset of cystic fibrosis. PLoS ONE 7, 17–19 (2012).

32. Loo, M. A. et al. Perturbation of Hsp90 interaction with nascent CFTR prevents its maturation and accelerates its degradation by the proteasome. EMBO J. 17, 6879–6887 (1998).

33. Loureiro, C. A. et al. A molecular switch in the scaffold NHERF1 enables misfolded CFTR to evade the peripheral quality control checkpoint. Sci. Signal. 8, ra48–ra48 (2015).

34. Hutt, D. M. et al. Silencing of the Hsp70-specific nucleotide-exchange factor BAG3 corrects the F508del-CFTR variant by restoring autophagy. J. Biol. Chem. 293, 13682–13695 (2018).

35. Sondo, E. et al. Pharmacological Inhibition of the Ubiquitin Ligase RNF5 Rescues F508del-CFTR in Cystic Fibrosis Airway Epithelia. Cell Chem. Biol. 25, 891–905.e8 (2018).

36. El Khouri, E., Le Pavec, G., Toledano, M. B. & Delaunay-Moisan, A. RNF185 is a novel E3 ligase of endoplasmic reticulum-associated degradation (ERAD) that targets cystic fibrosis transmembrane conductance regulator (CFTR). J. Biol. Chem. 288, 31177–31191 (2013).

37. Shishido, H., Yoon, J. S., Yang, Z. & Skach, W. R. CFTR trafficking mutations disrupt cotranslational protein folding by targeting biosynthetic intermediates. Nat. Commun. 11, 4258–4258 (2020).

38. Farsad, K. et al. Generation of high curvature membranes mediated by direct endophilin bilayer interactions. J. Cell Biol. 155, 193–200 (2001).

39. McDonald, E. F. et al. Structural Comparative Modeling of Multi-Domain F508del CFTR. Biomolecules 12, 471 (2022).

40. Zhang, Z., Liu, F. & Chen, J. Molecular structure of the ATP-bound, phosphorylated human CFTR. Proc. Natl. Acad. Sci. 115, 12757–12762 (2018).

41. Liu, F., Zhang, Z., Csanády, L., Gadsby, D. C. & Chen, J. Molecular Structure of the Human CFTR Ion Channel. Cell 169, 85–92 (2017).

42. Wang, R. Y.-R. et al. Automated structure refinement of macromolecular assemblies from cryo-EM maps using Rosetta. eLife 5, e17219 (2016).

43. He, L. et al. Restoration of domain folding and interdomain assembly by second site suppressors of the ΔF508 mutation in CFTR. FASEB J. 24, 3103–3112 (2010).

44. Lewis, H. A. et al. Structure and dynamics of NBD1 from CFTR characterized using crystallography and hydrogen/deuterium exchange mass spectrometry. J. Mol. Biol. 396, 406–430 (2010).

45. Yang, Z. et al. Structural stability of purified human CFTR is systematically improved by mutations in nucleotide binding domain 1. Biochim. Biophys. Acta - Biomembr. 1860, 1193–1204 (2018).

46. Dawson, J. E., Farber, P. J. & Forman-Kay, J. D. Allosteric Coupling between the Intracellular Coupling Helix 4 and Regulatory Sites of the First Nucleotide-binding Domain of CFTR. PLoS ONE 8, (2013).

47. McDonald, E. F., Oliver, K. E., Schlebach, J. P., Meiler, J. & Plate, L. Benchmarking AlphaMissense

48. Carolyn Padoa, Andrea Goldman, Trefor Jenkins, & Michele Ramsay. Cystic fibrosis carrier frequencies in populations of African origin. J. Med. Genet. 36, 41 (1999).

49. Zeiger, A. M. et al. Identification of CFTR variants in Latino patients with cystic fibrosis from the Dominican Republic and Puerto Rico. Pediatr. Pulmonol. 55, 533–540 (2020).

50. Farinha, C. M. et al. Revertants, low temperature, and correctors reveal the mechanism of F508del-CFTR rescue by VX-809 and suggest multiple agents for full correction. Chem. Biol. 20, 943–955 (2013).

51. He, L. et al. DNAJB12 and Hsp70 Triage Arrested Intermediates of N1303K-CFTR for ER Associated-Autophagy Lihua. Mol. Biol. Cell 44, 1689–1699 (2021).

52. Mendoza, J. L. et al. Requirements for efficient correction of Δf508 CFTR revealed by analyses of evolved sequences. Cell 148, 164–174 (2012).

53. Hurlbut, G. et al. EPS3.10 Novel CFTR modulator combinations directly address the ΔF508-CFTR NBD1 stability defect and enable full CFTR correction. J. Cyst. Fibros. 22, S45 (2023).

54. Kim, M. & Plate, L. Cystic Fibrosis Modulator Therapies: Bridging Insights from CF to other Membrane Protein Misfolding Diseases. Isr. J. Chem. e202300152 (2024) doi:10.1002/ijch.202300152.

55. Sabusap, C. M. et al. Analysis of cystic fibrosis–associated P67L CFTR illustrates barriers to personalized therapeutics for orphan diseases. JCI Insight 1, 1–10 (2016).

56. Pankow, S., Bamberger, C., Calzolari, D., Bamberger, A. & Yates III, J. R. Deep interactome profiling of membrane proteins by co-interacting protein identification technology. Nat. Protoc. 11, 2515–2515 (2016).

57. Welzel, T., Radtke, I., Meyer-Zaika, W., Heumann, R. & Epple, M. Transfection of cells with custom-made calcium phosphate nanoparticles coated with DNA. J. Mater. Chem. 14, 2213 (2004).

58. Mendenhall, J., Brown, B. P., Kothiwale, S. & Meiler, J. BCL::Conf: Improved Open-Source Knowledge-Based Conformation Sampling Using the Crystallography Open Database. J. Chem. Inf. Model. 61, 189–201 (2021).

59. Lemmon, G. & Meiler, J. Rosetta Ligand Docking with Flexible XML Protocols. in Computational Drug Discovery and Design (ed. Baron, R.) vol. 819 143–155 (Springer New York, New York, NY, 2012).

60. Song, Y. et al. High-Resolution Comparative Modeling with RosettaCM. Structure 21, 1735–1742 (2013).

