## Supplementary Information for "Proteostasis Landscapes of Cystic Fibrosis Variants Reveals Drug Response Vulnerability"

**for**

### Table of Contents

Supplementary Table S1………………………………………………………………………………..2

Supplementary Figures S1-S13…...……………………………………………………………..…..2-22

References……………….……………………………………………………………...……..23

.

**Supplemental Table S1. Allele frequencies of poorly responsive CFTR variants[1].**

| Allele | # of Patients | w/ F508del | w/o F508del |
|---|---|---|---|
| V520F | 155 | 105 | 50 |
| L558S | 33 | 19 | 14 |
| A559T | 85 | 39 | 46 |
| R560T | 340 | 243 | 97 |
| A561E | 15 | 7 | 8 |
| Y569D | 28 | 2 | 26 |
| R1066C | 211 | 124 | 87 |
| Total | 867 | | **328** |

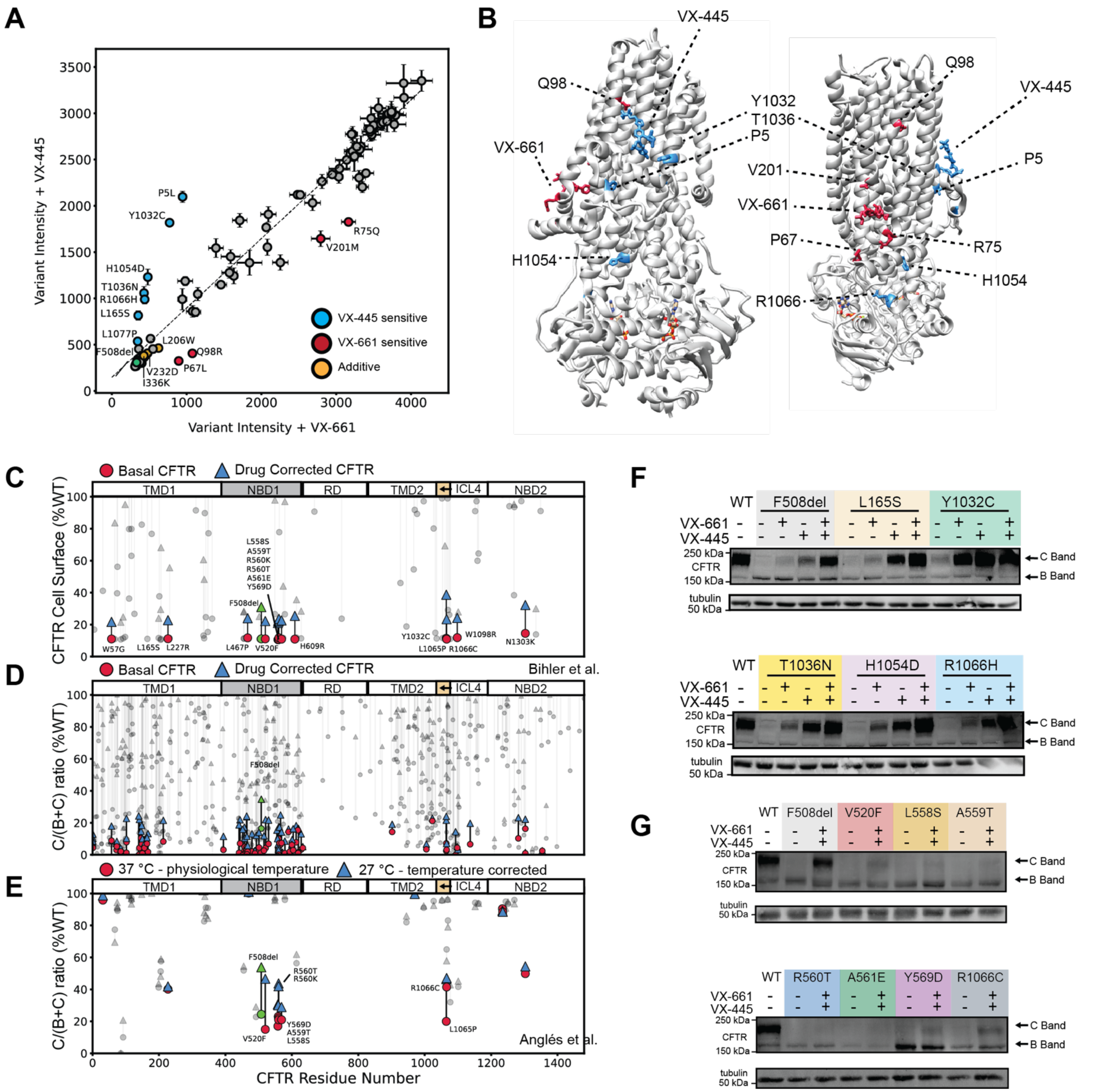


**Supplementary Figure S1. Selectively and poorly responsive CFTR variants with VX-661 and VX-445.**
**A.** Adapted from Figure 3B in[2] CFTR cell surface immune staining intensity comparing treatment with VX-661 versus VX-445 correctors with the best fit dotted line representing equivalent response to both correctors. Variants below the best-fit dotted line were selectively responsive to VX-661, while variants above the dotted line were selectively responsive to VX-445. Selectively sensitive variants annotated. Error bars represent standard deviations.
**B.** CFTR structure (PDB ID 8EIQ)[3] showing VX-445 sensitive variants proximate to the VX-445 binding site (blue) as well as VX-661 sensitive variants proximate to the VX-661 binding site (red).
**C.** CFTR cell surface immunostaining as assessed by deep mutational scanning (DMS) from[2]. CFTR residues are resolved by sequence position on the x-axis. Cell surface staining intensity on the y-axis is expressed as

percent WT (%WT) for basal (red circles) and dual corrector (VX-661 and VX-445) treatment (blue triangle) conditions. The black line connecting each variant basal level to the drug level represents the magnitude of drug response. For reference, approved variants are shown in grey. Variants with >100% WT drug response were removed for clarity. Unapproved, poorly responsive CFTR mutations cluster in the alpha-helical subdomain of NBD1 (residues 460-600) and inter-cellular loop4 (residues 1030-1080).

**D.** Reanalysis of CFTR trafficking data from[4]. Variants are resolved based on sequence position on the x-axis. The y-axis shows the measured C/B band ratio in percent WT (%WT) under basal (red circle) and dual corrector (VX-661 and VX-445) treatment (blue triangle) conditions. Approved variants are shown in grey. Variants with >25% WT drug response were removed for clarity.

**E.** Reanalysis of CFTR trafficking data from[5]. The Y-axis shows measured C/(C+B) band ratio representing the trafficking efficiency in percent WT (%WT) under standard temperature (37 C – red circles) and reduced temperature (27 C – blue triangles) conditions with a black line representing temperature response magnitude. FDA-approved variants were filtered out and are shown in grey.

**F.** Representative Western blots depicting VX-445 selectively responsive variants L165S, Y1032C, T1036N, H1054D, and R1066H CFTR under basal, VX-661, VX-445, and dual drug treatment conditions. WT and F508del CFTR are shown as controls.

**G.** Western blots depicting poorly responsive variants V520F, L558S, A559T, R560T, A561E, Y569D, and R1066C CFTR under basal versus VX-661 + VX-445 dual treatment conditions. WT and F508del CFTR are shown as controls. The lack of maturely glycosylated C band CFTR during dual drug treatment indicates these variants are poorly responsive in the transient HEK293T cell system.

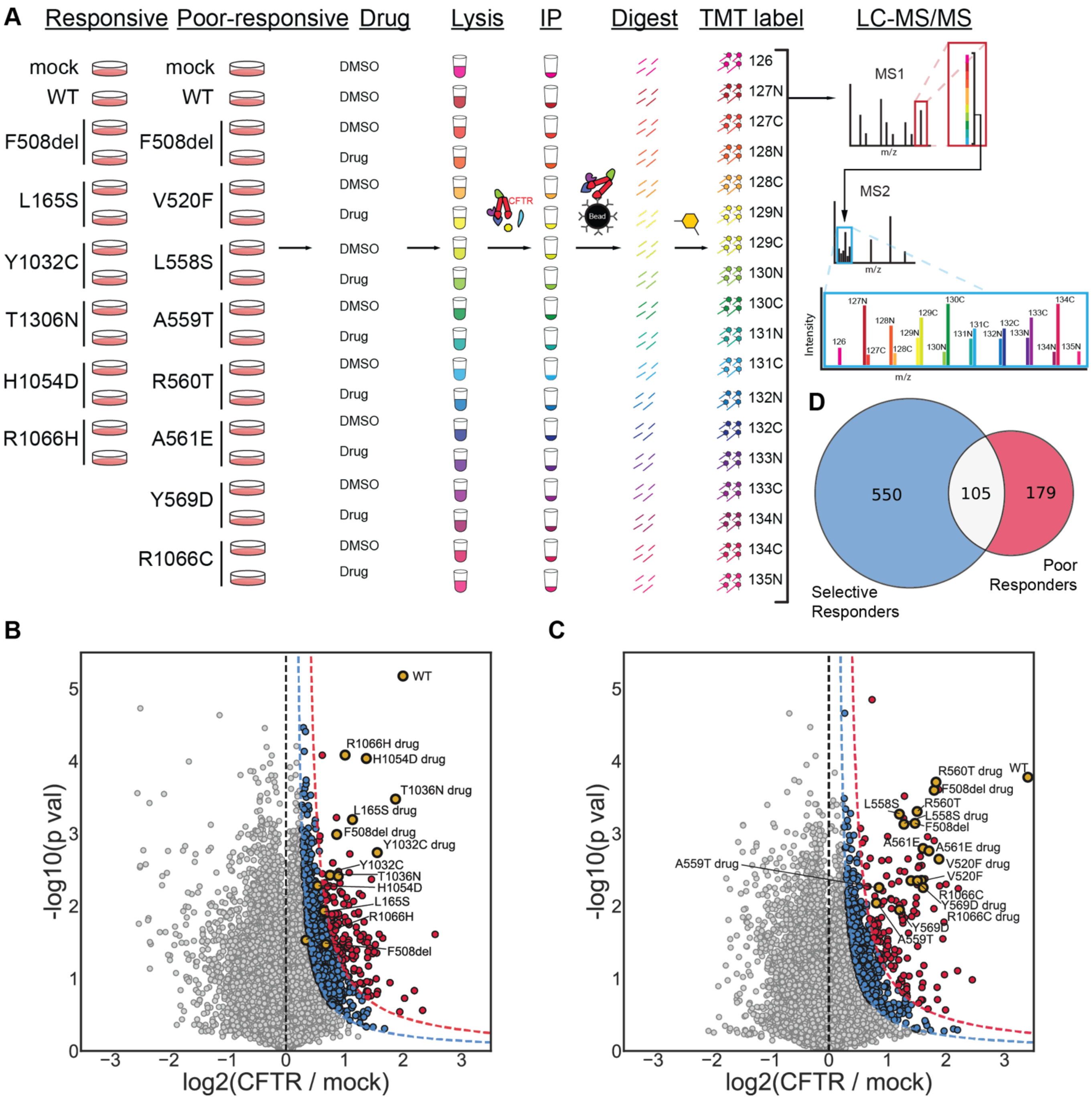


**Supplemental Figure S2. Quantitative interactomics of VX-445 selectively and poorly responsive variants.** **A.** Quantitative affinity purification mass spectrometry interactomics workflow for identifying CFTR interactors. HEK293T cells are transiently transfected with CFTR variants of interest in a set of either VX-445 selective responders or a set of poorly responsive variants. Cells are then drug-treated with CFTR correctors, lysed, collected, and normalized. CFTR interactors are immunoprecipitated, eluted, and digested into their component peptides, which are labeled with isobaric TMT labels for multiplexed mass spectrometry analysis and quantification.

**B.** Volcano plot of VX-445 selective responder interactors. Interactors were identified using a curved cutoff at 0.5σ for medium confidence (blue) and 1σ for high confidence (red) proteins. CFTR, shown in gold, of each variant was enriched across conditions. Interacting proteins across selectively responsive variants were added to a master list comprising 655 interactors for this data set.

**C.** Volcano plot of poorly responsive interactors identified using a curved cutoff at 0.5σ for medium confidence (blue) and 1σ for high confidence (red) proteins. CFTR, shown in gold, of each variant was enriched for all mutants with and without drug treatment. Interacting proteins across poorly responsive variants were added to a master list comprising 284 interactors for this data set.

**D.** Overlap of 105 shared interactors between the 655 interactors identified in the VX-445 selectively responsive dataset and 284 interactors identified in the poor-responder dataset.

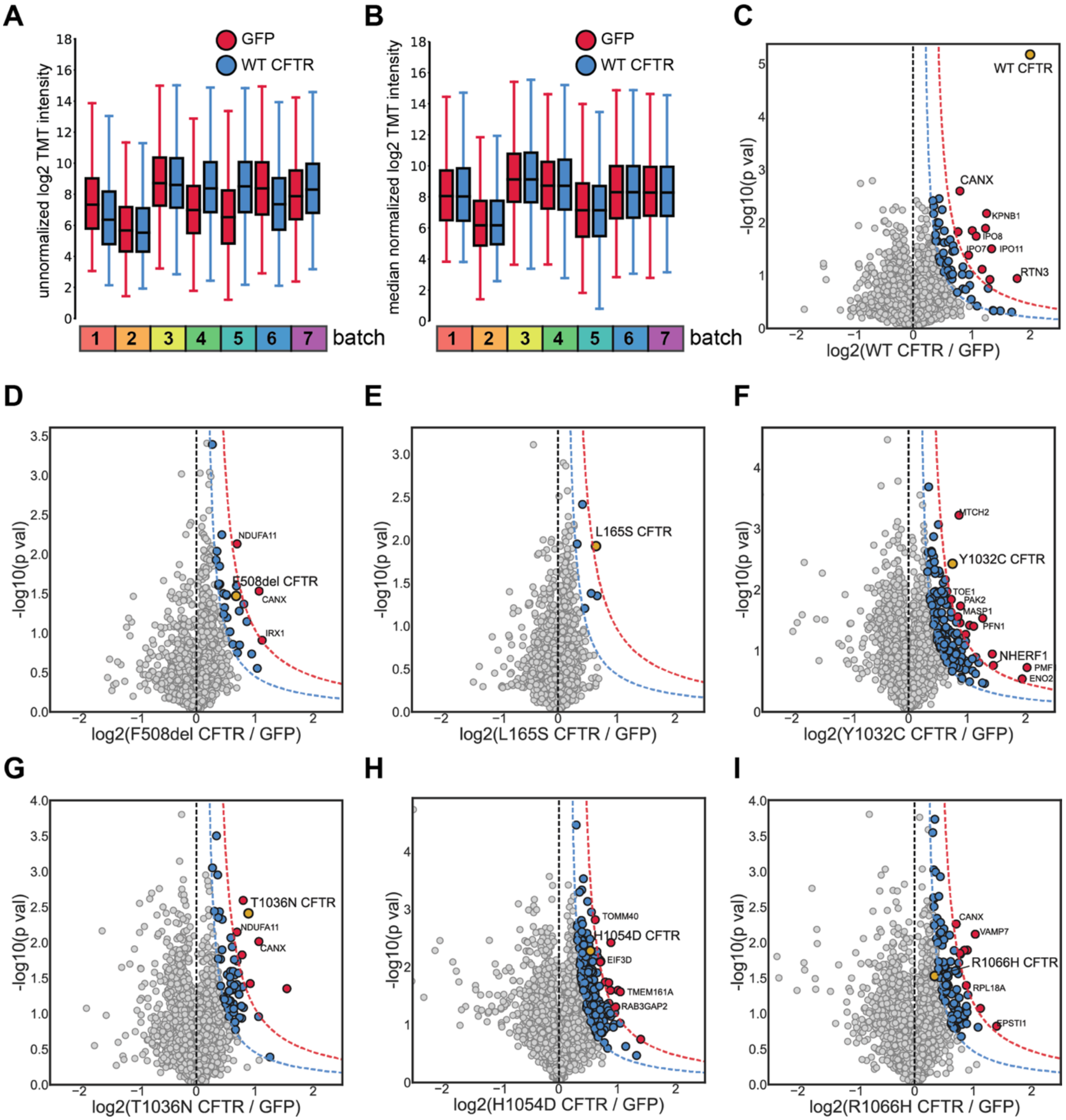


**Supplemental Figure S3. Identification of statistically significant interactors with VX-445 responsive CFTR variants under basal (DMSO) conditions.**

**A.** Box and whisker plot showing the unnormalized TMT channel intensity distributions of GFP mock control and WT CFTR for all 7 LC-MS/MS runs included in the selectively responsive variants dataset. The box limits represent the upper and lower quartiles with a line at the median; the whiskers represent the limits 1.5 times the interquartile range.

**B.** Box and whisker plot showing the corresponding median normalized TMT channel intensity distributions of GFP mock control and WT CFTR for the 7 LC-MS/MS run. Median normalization of TMT quantification for a given run results int the median of each run for different conditions, e.g. GFP and WT shown here, approximately equal. Box plots represented the same as in A.

**C.** Volcano plot of WT interactors. Interactors were identified using a curved cutoff at 0.5σ for medium confidence (blue) and 1σ for high confidence (red) proteins; CFTR shown in gold. Interacting proteins across selectively responsive variants were added to a master list comprising 655 interactors for this data set.
**D.** Volcano plot of F508del + DMSO interactors.
**E.-I.** Volcano plot of selectively responsive variants + DMSO interactors for L165S, Y1032C, T1036N, H1054D, R1066H respectively. L165S showed little interactor enrichment. Y1032C shows enrichment of previously identified and characterized CFTR interactor NHERF1, a plasma membrane (PM) scaffolding protein important for CFTR peripheral stability and quality control[6]. T1036N and R1066H show enrichment of previously identified and characterized CFTR interactor CANX[7,8].

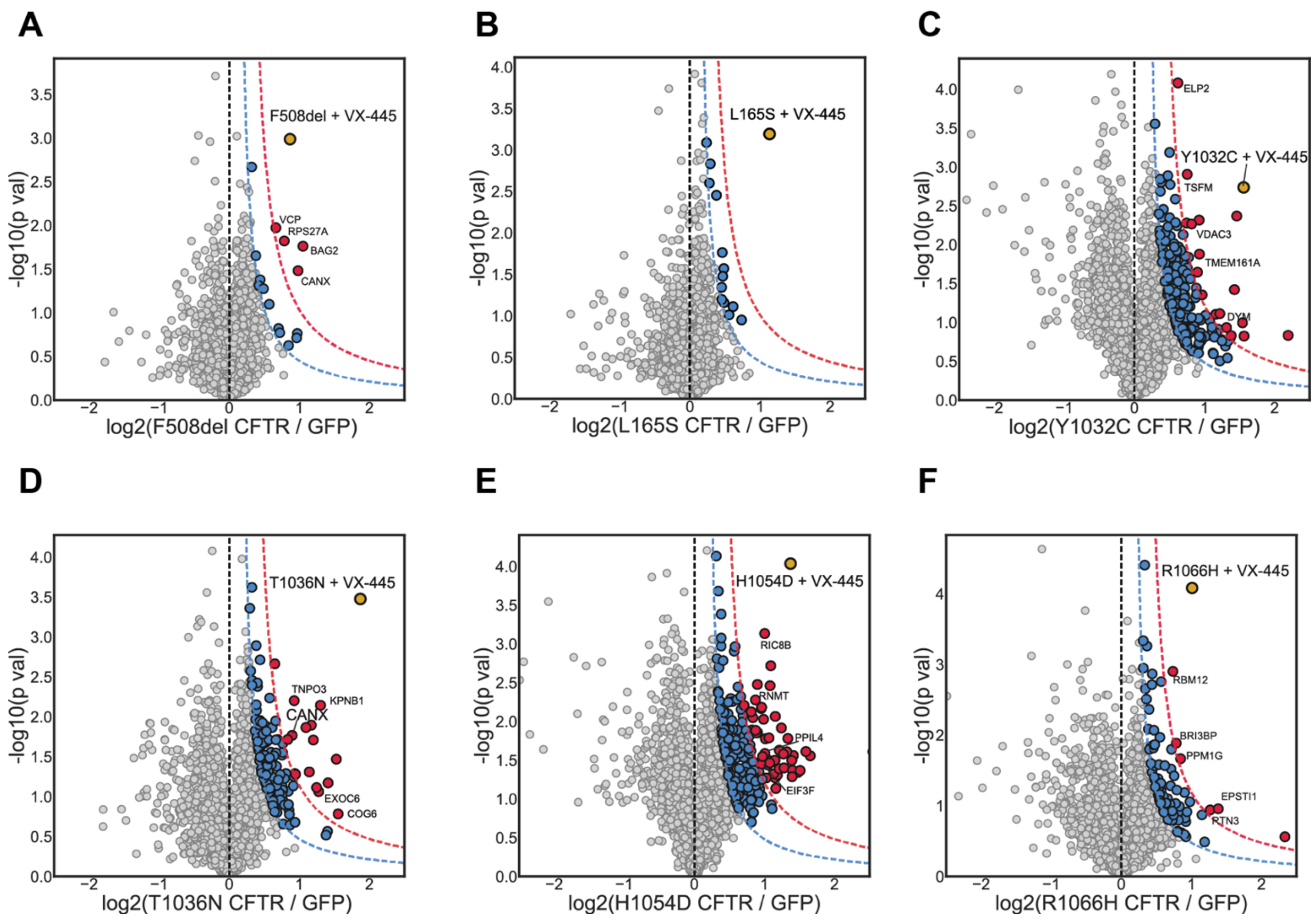


**Supplemental Figure S4. Identification of statistically significant interactors with VX-445 responsive CFTR variants under drug treatment (3 µM VX-445) conditions.**

**A.** Volcano plot of F508del + VX-445 interactors. Interactors were identified using a curved cutoff at 0.5σ for medium confidence (blue) and 1σ for high confidence (red) proteins, CFTR shown in gold. Interacting proteins across selectively responsive variants were added to a master list comprising 655 interactors for this data set.

**B.-F.** Volcano plot of selectively responsive variants + VX-445 interactors for L165S, Y1032C, T1036N, H1054D, R1066H respectively. Colored as in **A.**

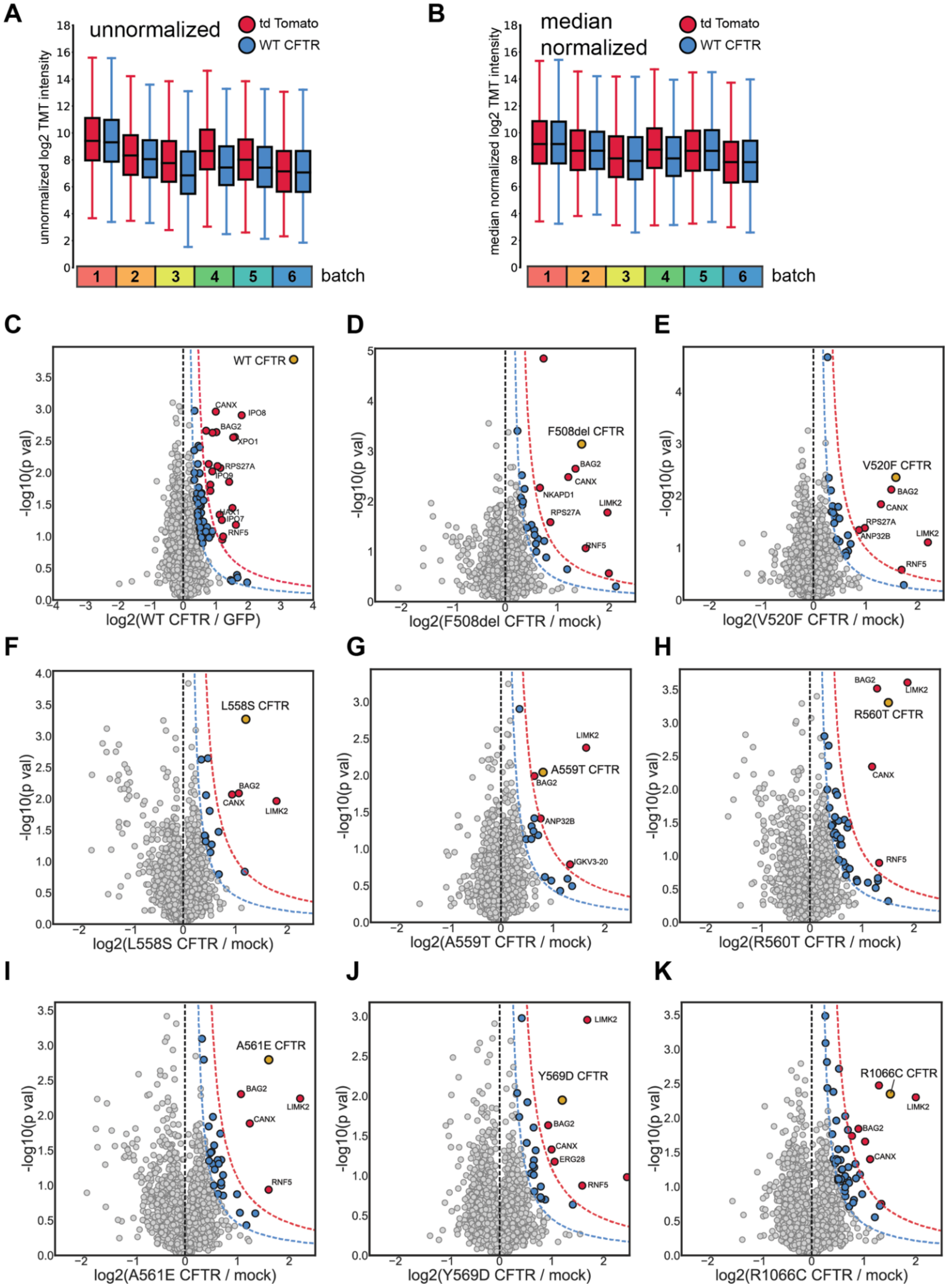
A
unnormalized
td Tomato
WT CFTR
unnormalized log2 TMT intensity
batch
B
median normalized
median normalized log2 TMT intensity
batch
C
WT CFTR
CANX
IPO8
BAG2
XPO1
RPS27A
IPO9
HAX1
IPO7
RNF5
-log10(p val)
log2(WT CFTR / GFP)
D
F508del CFTR
BAG2
CANX
NKAPD1
RPS27A
LIMK2
RNF5
log2(F508del CFTR / mock)
E
V520F CFTR
BAG2
CANX
RPS27A
ANP32B
LIMK2
RNF5
log2(V520F CFTR / mock)
F
L558S CFTR
BAG2
CANX
LIMK2
log2(L558S CFTR / mock)
G
LIMK2
A559T CFTR
BAG2
ANP32B
IGKV3-20
log2(A559T CFTR / mock)
H
BAG2
LIMK2
R560T CFTR
CANX
RNF5
log2(R560T CFTR / mock)
I
A561E CFTR
BAG2
LIMK2
CANX
RNF5
log2(A561E CFTR / mock)
J
LIMK2
Y569D CFTR
BAG2
CANX
ERG28
RNF5
log2(Y569D CFTR / mock)
K
R1066C CFTR
LIMK2
BAG2
CANX
log2(R1066C CFTR / mock)

**Supplemental Figure S5. Identification of statistically significant interactors with poorly-responsive CFTR variants under basal conditions.**

**A.** Box and whisker plot showing the unnormalized TMT channel intensity distributions of tdTomato mock control and WT CFTR for all 6 LC-MS/MS runs included in the selectively responsive variants dataset. The box limits represent the upper and lower quartiles with a line at the median; the whiskers represent the limits 1.5 times the interquartile range.

**B.** Box and whisker plot showing the median normalized TMT channel intensity distributions of td-Tomato mock control and WT CFTR for the corresponding 6 LC-MS/MS run. Median normalization of TMT quantification for a given run results int the median of each run for different conditions, e.g. td-Tomato and WT shown here, approximately equal. Box plots represented the same as in A.

**C.** Volcano plot of WT CFTR interactors. Interactors were identified using a curved cutoff at 0.5σ for medium confidence (blue) and 1σ for high confidence (red) proteins, CFTR shown in gold. Interacting proteins across poorly responsive variants and controls were added to a master list comprising 284 interactors for this data set.

**D.** Volcano plot of F508del CFTR + DMSO interactors. Colored as in **C.**

**E.-K.** Volcano plots of poorly responsive CFTR variants + DMSO interactors for V520F, L558S, A559T, R560T, A561E, Y569D, and R1066C, respectively. Colored as in **C.** CANX[7,8], BAG2[9,10], and RNF5[11], as well as new interactor RPS27A, a small ribosomal subunit protein involved in ribosomal quality control[12,13], and LIMK2, and serine kinase implicated in actin filament assembly and Rho GTPase signaling[14,15] Interestingly, interaction with CANX[3], BAG2, RNF5, and LIMK2 were remarkably consistent amongst poorly responsive variants.

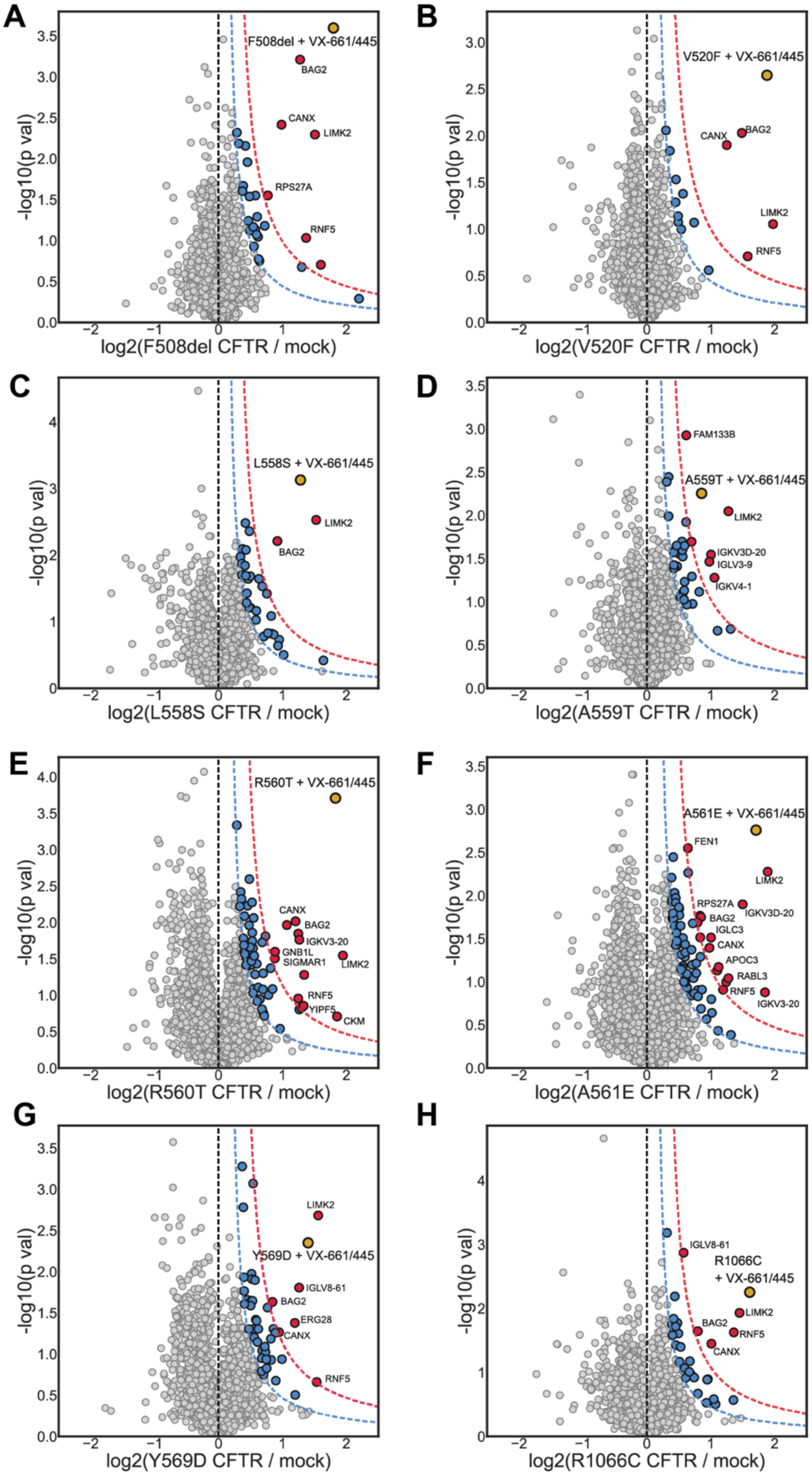


**Supplemental Figure S6. Identification of statistically significant interactors with poorly responsive CFTR variants under drug treatment (3 µM VX-661 + 3 µM VX-445) conditions.**

**A.** Volcano plot of F508del CFTR + 3 µM VX-661 + 3 µM VX-445 interactors. Interactors were identified using a curved cutoff at 0.5σ for medium confidence (blue) and 1σ for high confidence (red) proteins, CFTR shown in gold.

**B-H.** Volcano plots of poorly responsive CFTR variants + 3 µM VX-661 + 3 µM VX-445 interactors for V520F, L558S, A559T, R560T, A561E, Y569D, and R1066C, respectively. Colored as in **A.**

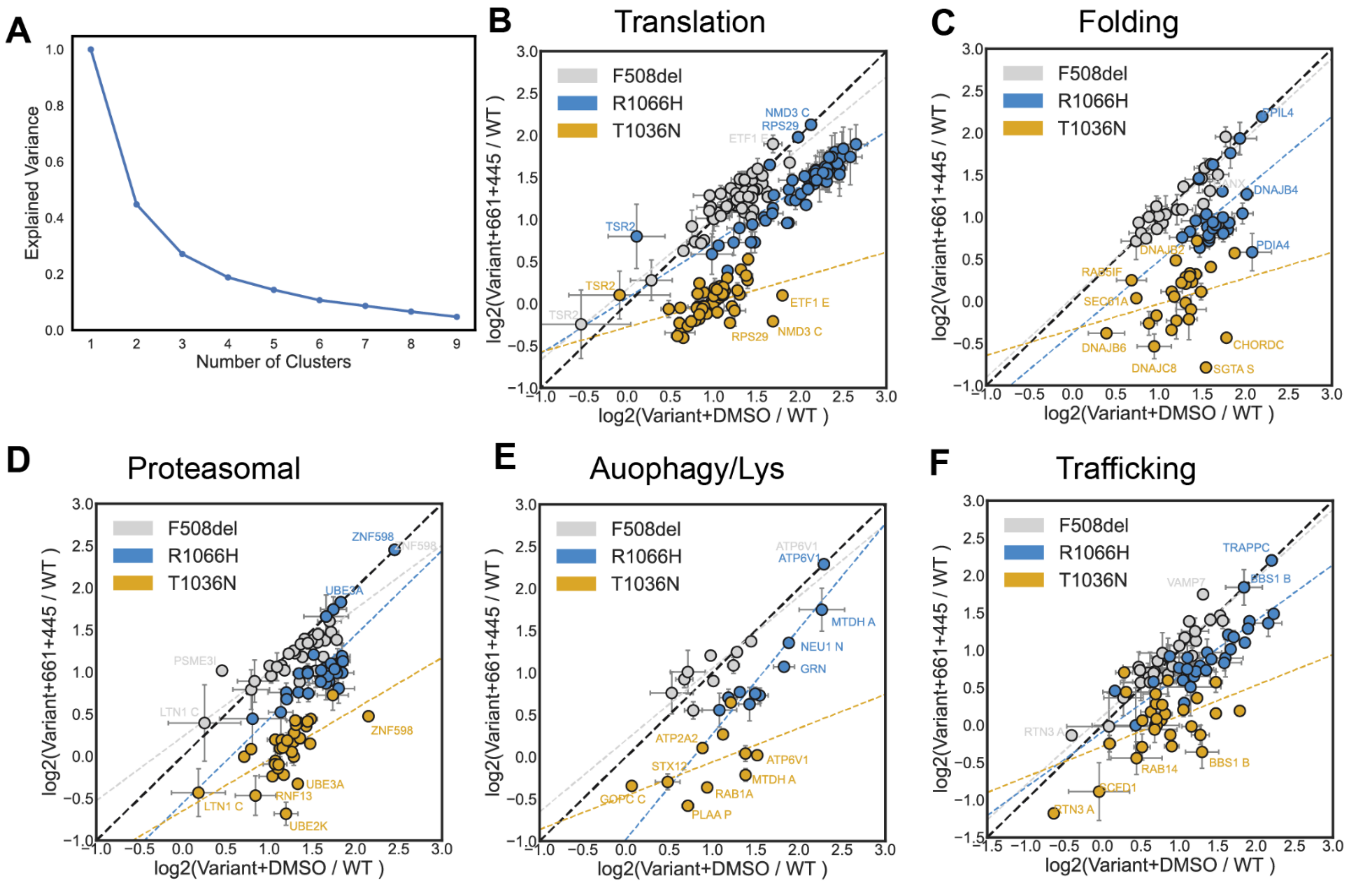


**Supplemental Figure S7. Correlation between DMSO and 3 µM VX-445 protein interactor quantification levels organized by pathway for VX-445 selectively responsive CFTR variants.**

**A.** Elbow plot of K means clustering cluster number versus explained variance for 655 interactors across all selectively responsive variants. The inflection point of the curve (elbow) occurs between 4 and 6 clusters; hence, we chose 4 clusters to cluster the Spearman rank order correlation heatmap in **Figure 2A**.

**B.-F.** Individual protein quantification correlation between DMSO and VX-445 treatment for F508del (grey), T1036N (gold), and R1066H (blue) for proteostasis pathways. A normal line (black dotted line) represents x=y or no change with the drug. A least squared linear best fit dotted line colored for each variant represented the off-normal axis behavior during drug treatment.

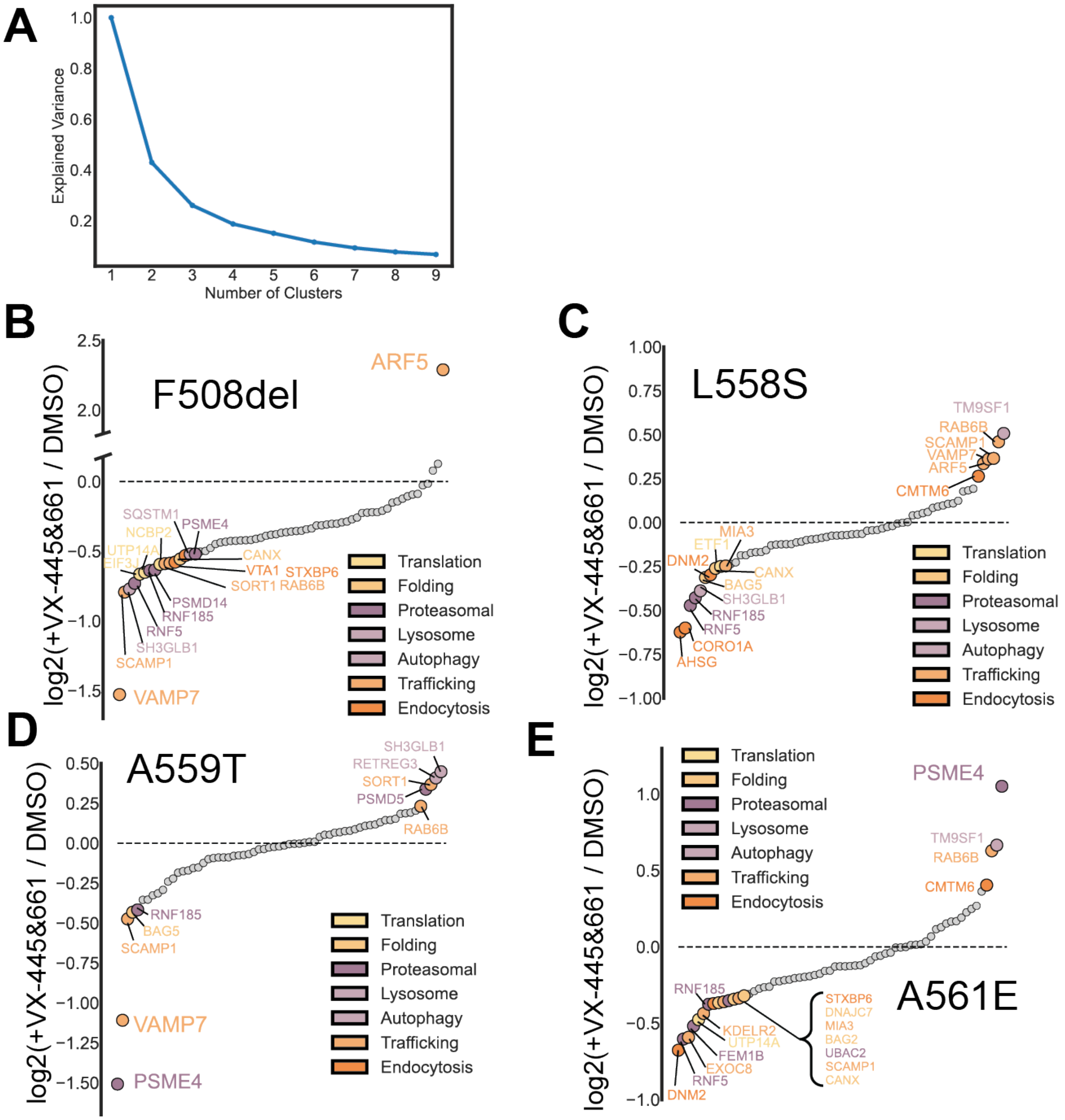


**Supplemental Figure S8. Clustering and waterfall plots of poorly responsive variants.**

**A.** Elbow plot of K means clustering cluster number versus explained variance for 284 interactors across all selectively responsive variants. The curve's inflection point (elbow) occurs between 5 and 7 clusters; hence, we chose 5 clusters to cluster the Spearman rank order correlation heatmap in **Figure 3A**.

**B.** A waterfall plot of the log2 fold change between F508del + 3 µM VX-661/3 µM VX-445 over F508del + DMSO organized by fold change rank order. Interactors above zero are greater with drug treatment, and interactors under zero are greater in the basal state. Interactors greater than one-half a standard deviation of positive values are colored by pathway; likewise, for interactors less than one-half a standard deviation of negative values.

**C-E.** A waterfall plot of the log2 fold change between poorly responsive variants + 3 µM VX-661/3 µM VX-445 over respective variant + DMSO organized by fold change rank order and color by pathway as in **B**.

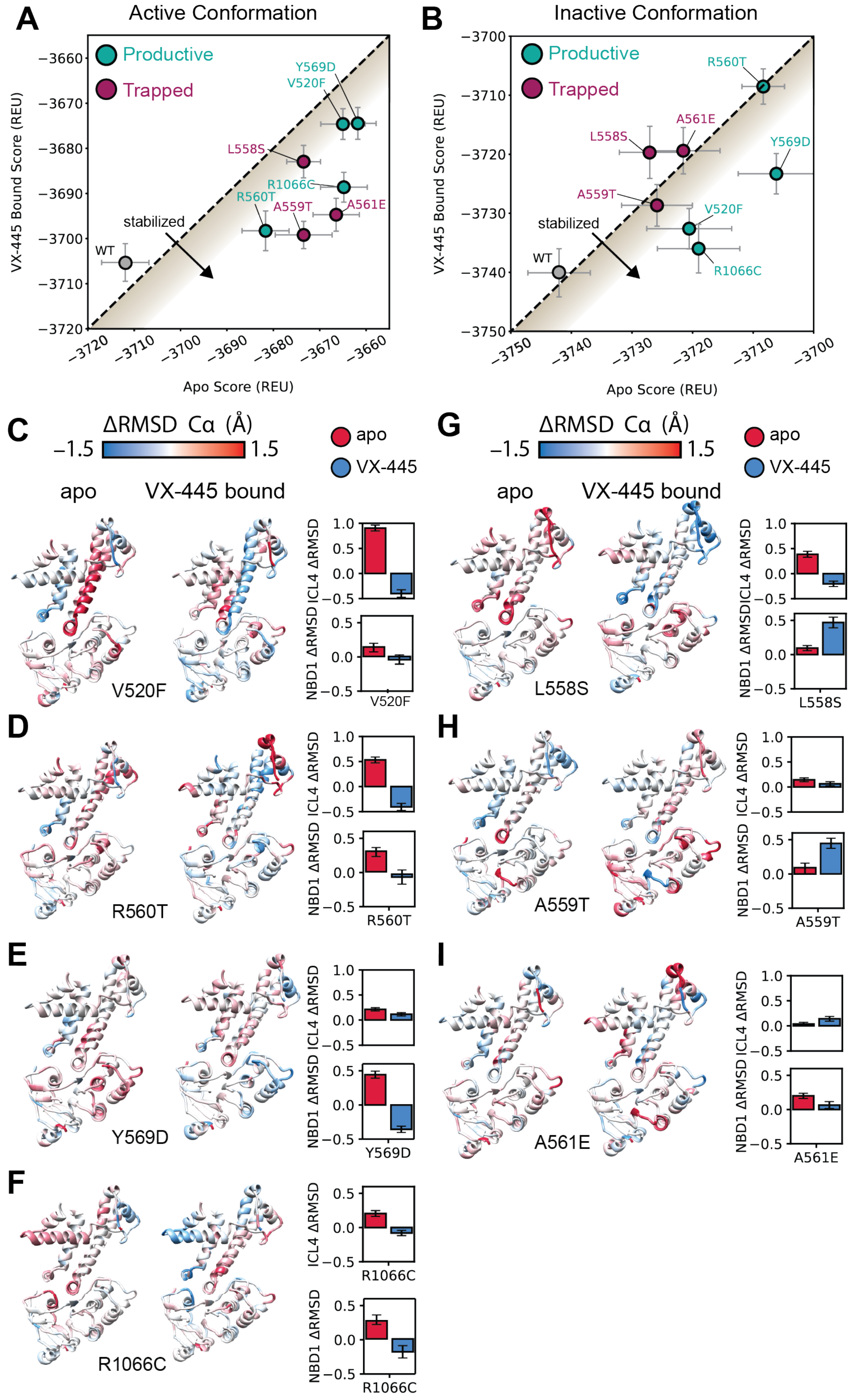
A
Active Conformation
Productive
Trapped
VX-445 Bound Score (REU)
Apo Score (REU)
Y569D
V520F
L558S
R1066C
R560T
A559T
A561E
stabilized
WT
B
Inactive Conformation
C
ΔRMSD Cα (Å)
−1.5
1.5
apo
VX-445
VX-445 bound
ICL4 ΔRMSD
NBD1 ΔRMSD
V520F
G
L558S
D
R560T
H
A559T
E
Y569D
I
A561E
F
R1066C

**Supplemental Figure S9. Rosetta scores and inactive structural modeling data in the apo and VX-445 bound state for poorly responsive variants.**

**A.** The average Rosetta energy scores of the active conformation VX-445-bound state are plotted against the apo state CFTR models for productive proteostasis poor responders (purple) and trapped proteostasis poor responders (green). A dotted black line with a slope of one intersecting the origin (no effect) is shown for reference. Points represent the average Rosetta energy score, and the error bars represent the standard error of the mean across the 100 lowest (10%) energy-scoring models from an ensemble of 1000 active state models generated for each variant. Variants falling below the dotted line representing x=y indicate thermodynamic stabilization by VX-445, whereas variants above the line are destabilized, and variants on the line remain unchanged.

**B.** The average Rosetta energy scores of the inactive conformation VX-445-bound state are plotted against the apo state CFTR models, colored as in **A**. Points represent the average Rosetta energy score, and the error bars represent the standard error of the mean.

**C.** Poorly responsive V520F ΔRMSD in the inactive conformation mapped onto CFTR structure, ICL1, ICL4, lasso motif, and NBD1 shown for clarity and ILC4 and NBD1 (residues 485-572) ΔRMSD in the inactive conformation quantification. The left structure labeled apo represents the ensemble RMSD in the inactive conformation of V520F (lowest 10% of models) minus the ensemble RMSD of WT CFTR (lowest 10% of models). The right structure labeled VX-445 bound represents the ensemble RMSD of V520F + VX-445 minus the ensemble RMSD of V520F apo. The bar graph shows the average, unnormalized ensemble RMSD for the apo and VX-445 bound V520F models.

**D-I.** Poorly responsive R560T apo ΔRMSD in the inactive conformation normalized to WT (left) and VX-445 bound ΔRMSD in the inactive conformation normalized to apo R560T mapped onto CFTR ICL1/4, lasso, and NBD1. ΔRMSD quantification for ICL4 and NBD1 (485-572) is shown as a bar graph. **C.** Y656D ΔRMSD. **D.** R1066C ΔRMSD. **E.** L558S ΔRMSD. **F.** A559T ΔRMSD. **G.** A561E ΔRMSD.

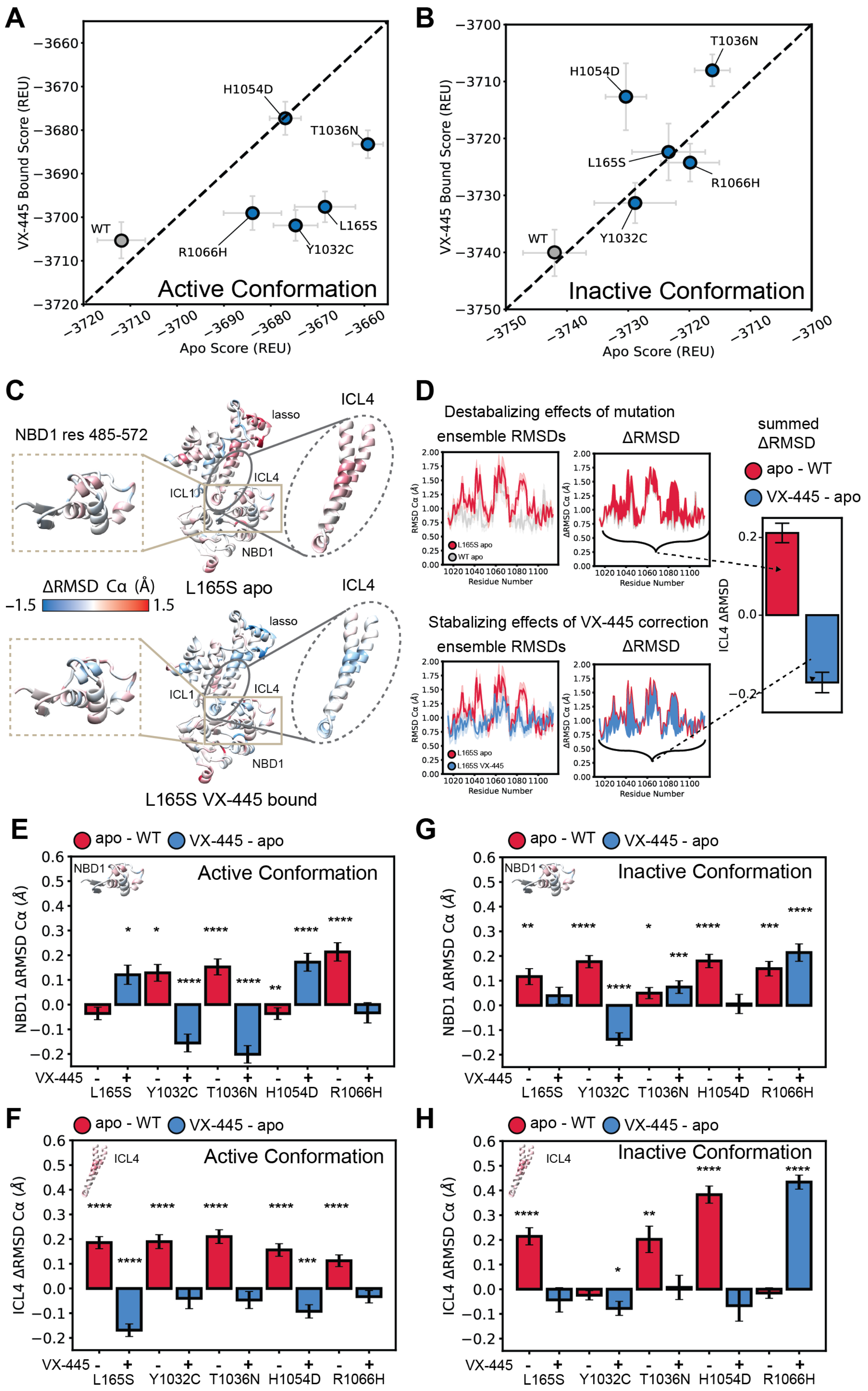
A
Active Conformation
VX-445 Bound Score (REU)
Apo Score (REU)
WT
H1054D
T1036N
R1066H
L165S
Y1032C
B
Inactive Conformation
VX-445 Bound Score (REU)
Apo Score (REU)
T1036N
H1054D
L165S
R1066H
Y1032C
WT
C
NBD1 res 485-572
lasso
ICL4
ICL1
NBD1
ΔRMSD Cα (Å)
−1.5
1.5
L165S apo
L165S VX-445 bound
D
Destabalizing effects of mutation
ensemble RMSDs
ΔRMSD
summed ΔRMSD
apo - WT
VX-445 - apo
ICL4 ΔRMSD
L165S apo
WT apo
Residue Number
Stabalizing effects of VX-445 correction
L165S VX-445
E
apo - WT
VX-445 - apo
Active Conformation
NBD1 ΔRMSD Cα (Å)
VX-445
L165S
Y1032C
T1036N
H1054D
R1066H
G
Inactive Conformation
F
ICL4 ΔRMSD Cα (Å)
Active Conformation
H
Inactive Conformation

**Supplemental Figure S10. RosettaCM bound to VX-445 versus apo state reveals the structural basis of VX-445 selective response.**

**A.** The average Rosetta energy scores of the active conformation VX-445-bound state are plotted against the apo state CFTR models for responsive variants considered in this study. A dotted black line with a slope of one intersecting the origin (no effect) is shown for reference. Points represent the average Rosetta energy score, and the error bars represent the standard error of the mean across the 100 lowest (10%) energy-scoring models from an ensemble of 1000 active state models generated for each variant. VX-445 stabilized variants falling below the dotted line.

**B.** The average Rosetta energy scores of the inactive conformation VX-445-bound state are plotted against the apo state selectively responsive CFTR variants models. Points represent the average Rosetta energy score, and the error bars represent the standard error of the mean.

**C.** Comparison of variant instability and VX-445 conferred stability. On top, the ΔRMSD between apo L165S and WT CFTR in the active conformation was mapped onto PDB ID 6MSM[16]. On the bottom, the ΔRMSD between VX-445 bound L165S and apo L165S CFTR in the active conformation was mapped onto PDB ID 6MSM[16]. Intercellular Loop 1 (ICL1), ICL4, Lasso motif, and NBD1 are shown for clarity. NBD1 ΔRMSD (residues 485-572) were regarded because they contact TMD2 via ICL4, and ICL4 was also quantified.

**D.** Quantification of average ΔRMSD in ICL4 as an example calculation. On top, the ensemble RMSD by residue is shown for apo L165S and WT CFTR, followed by a plot showing the ΔRMSD between the RMSDs shaded in red. This represents the instability caused by the L165S mutation in ICL4. On the bottom, the ensemble RMSD by residue is shown for VX-445 bound L165S and apo L165S CFTR, followed by a plot showing the ΔRMSD between the RMSDs shaded in blue. This represents the stability conferred by VX-445 to L165S CFTR in ICL4. Both ΔRMSD values are summed in the quantification bar plot shown at the end and in **E-H** below.

**E.** ΔRMSD of NBD1 residues 485-572 for selectively responsive variants in the active conformation. Error bars represent the standard error of the mean, and colored as in **D.** Statistical significance was calculated using a non-parametric Wilcoxon signed-rank test compared to zero to determine if distributions were significantly different from zero, and p values were depicted by *< 0.05, **< 0.01, ***< 0.001, and ****< 0.0001.

**F.** ΔRMSD of ICL4 for selectively responsive variants in the active conformation. High ΔRMSD changes suggest VX-445 confers substantial stability to ICL4 in the active conformation. Error bars represent the standard error of the mean, and colored as in **D.**

**G.** ΔRMSD of NBD1 residues 485-572 for selectively responsive variants in the inactive conformation. Error bars represent the standard error of the mean, and colored as in **D.**

**H.** ΔRMSD of ICL4 for selectively responsive variants in the inactive conformation. Error bars represent the standard error of the mean, and colored as in **D.**

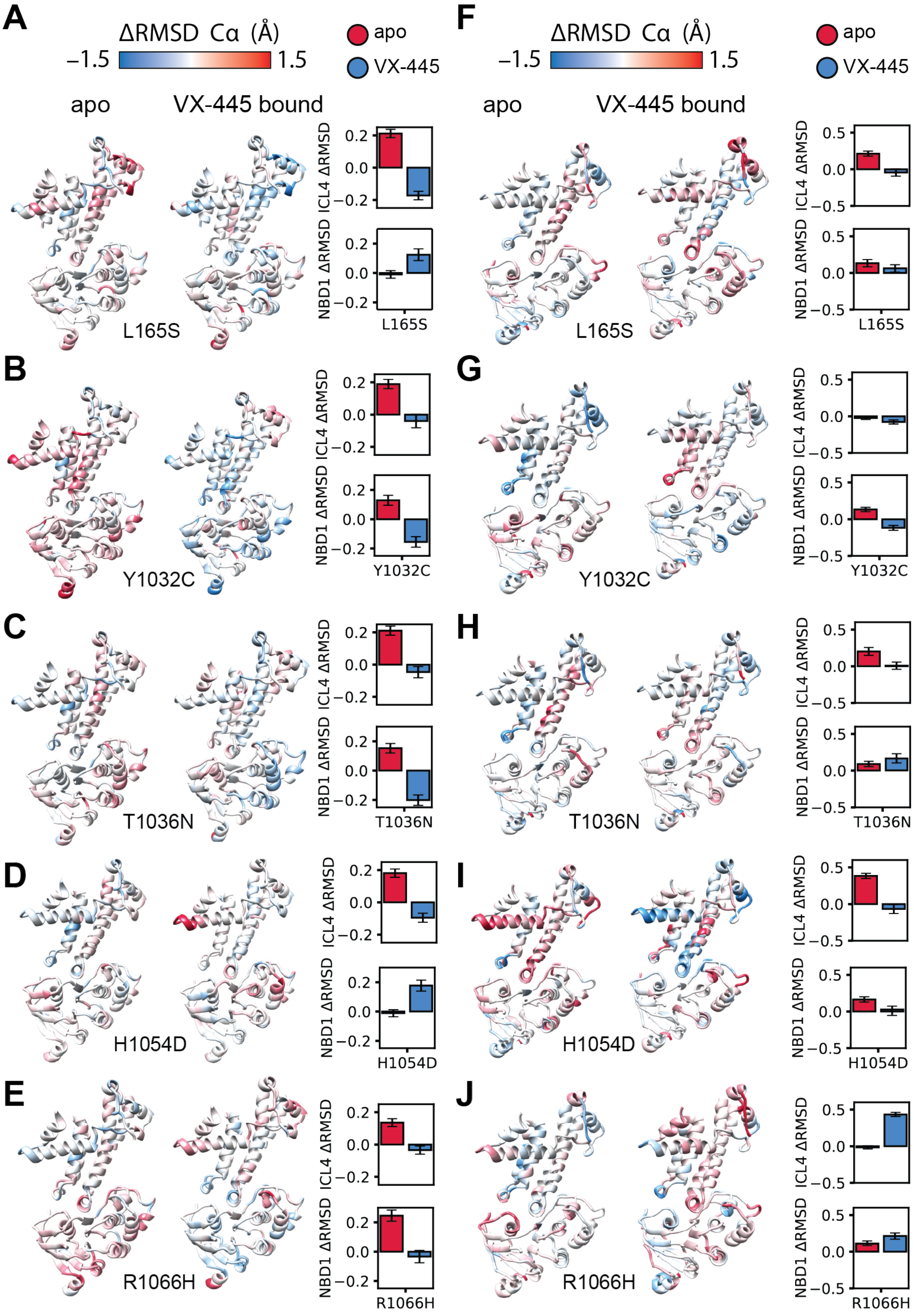


**Supplemental Figure S11. Active and inactive state structural modeling data of the apo and VX-445 bound selectively responsive variants.**

**A.** Selectively responsive L165S ΔRMSD in the active conformation mapped onto CFTR structure 6MSM, ICL1, ICL4, Lasso motif, and NBD1. The left structure labeled apo represents the ensemble RMSD in the active conformation of L165S (lowest 10% of models) minus the ensemble RMSD of WT CFTR (lowest 10% of models). The right structure labeled VX-445 bound represents the ensemble RMSD of L165S + VX-445 minus the ensemble RMSD of L165S apo. The bar graph shows the ΔRMSD for ICL4 and NBD1 (residues 485-572) which contact TMD2.
**B-E.** Selectively responsive apo/WT ΔRMSD in the active conformation (left) and VX-445 bound/apo ΔRMSD in the active conformation mapped onto CFTR 6MSM ICL1/4, lasso, and NBD1. ΔRMSD quantification for ICL4 and NBD1 (residues 485-572) which contact TMD2. **B.** Active state Y1032C ΔRMSD. **C.** Active state T1036N ΔRMSD. **D.** Active state H1054D ΔRMSD. **E.** Active state R1066H ΔRMSD.
**F-J.** Selectively responsive apo/WT ΔRMSD in the inactive conformation (left) and VX-445 bound/apo ΔRMSD in the active conformation mapped onto CFTR 5uak ICL1/4, lasso, and NBD1. ΔRMSD quantification for ICL4 and NBD1 (residues 485-572) which contact TMD2. **F.** Inactive state L165S ΔRMSD. **G.** Inactive state Y1032C ΔRMSD. **H.** Inactive state T1036N ΔRMSD. **I.** Inactive state H1054D ΔRMSD. **J.** Inactive state R1066H ΔRMSD.

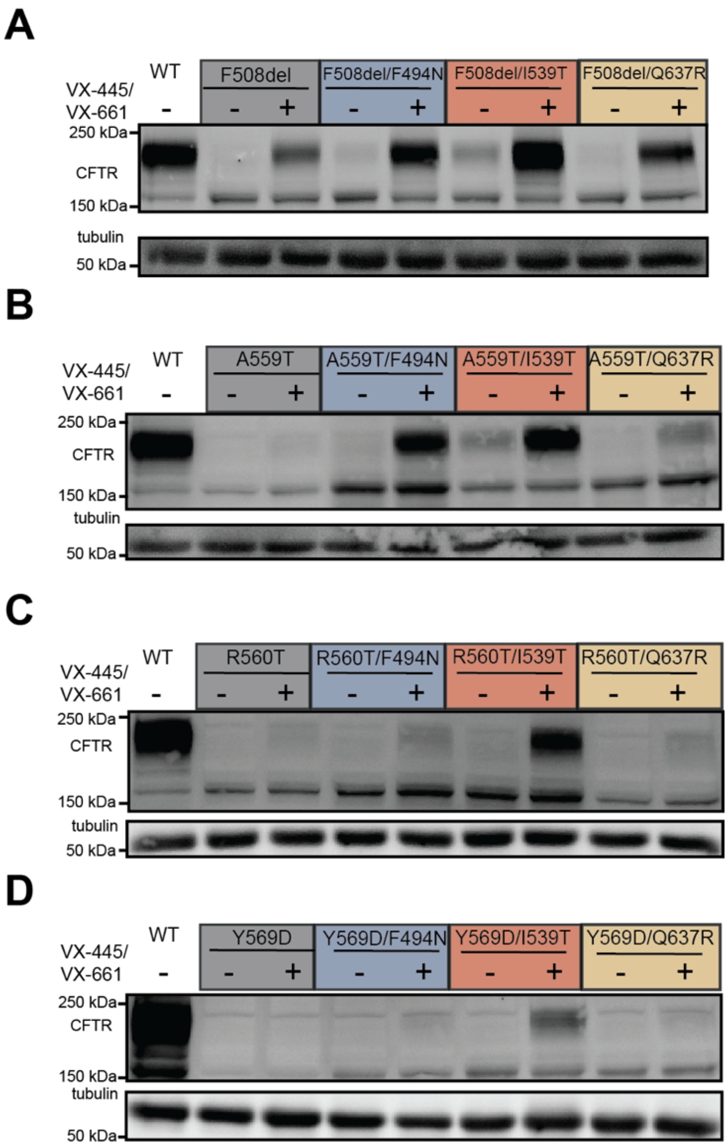


**Supplemental Figure S12. Representative western blots of secondary stabilizing mutation rescue of poorly responsive CFTR variants.**

**A.** Representative Western blots and quantifications depicting F508del CFTR under basal and dual drug treatment conditions combined with secondary stabilizing mutations F494N, I539T, and Q637R.

**B-D.** Representative western blots and blot quantifications depicting poorly responsive variants A559T, R560T, and Y569D CFTR under basal and dual drug treatment conditions in combination with secondary stabilizing mutations F494N, I539T, and Q637R. WT CFTR is shown as a control.

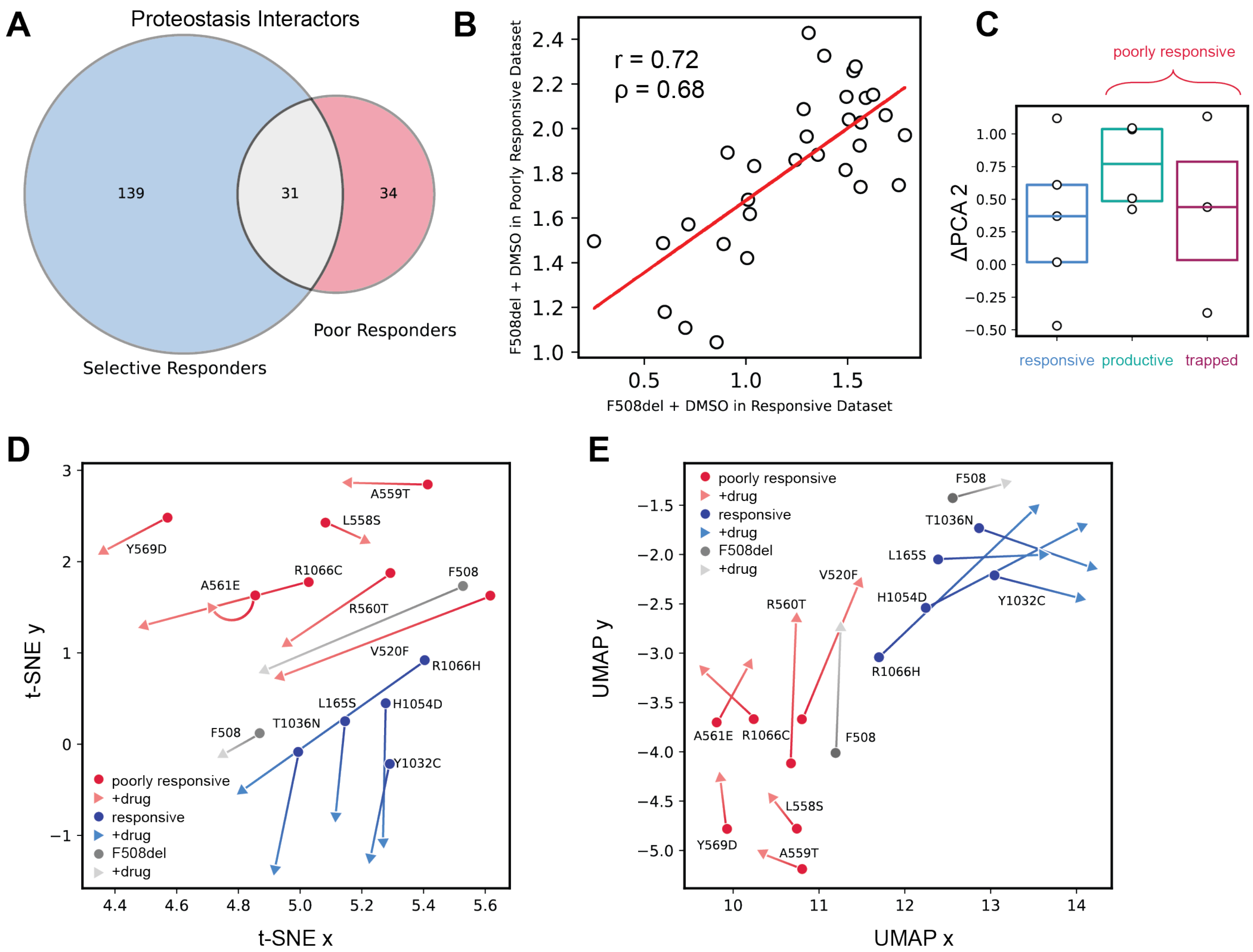


**Supplemental Figure S13. Comparing selectively and poorly responsive proteostasis interactors through overlap, correlation, and dimensionality reduction.**

**A.** Of the 170 selectively responsive proteostasis interactors (**Figure 2B**) and the 65 poorly responsive proteostasis interactors (**Figure 3B**). There is an overlap of 31 proteostasis proteins.

**B.** F508del + DMSO is the only condition found in both datasets. The correlation of the 31 overlapping proteostasis interactors has a Pearson correlation (r) of 0.72 and a Spearman correlation (ρ) of 0.68.

**C.** Distributions of the second principal component shift (ΔPC2).

**D.** t-distributed stochastic neighbor embedding (t-SNE) dimensionality reduction of 31 overlapping proteostasis interactors. Basal conditions are shown as circles, and drug conditions (VX-445 or VX-661+VX-445, respectively) are shown as triangles. A line connecting basal and drug points for the same variants shows how the correction shifts interactions for each variant.

**E.** Uniform Manifold Approximation (UMAP) dimensionality reduction of 31 overlapping proteostasis interactors. Colored as in **D.**

### References


1. The Clinical and Functional TRanslation of CFTR (CFTR2); available at http://cftr2.org.
2. McKee, A. G. *et al.* General trends in the effects of VX-661 and VX-445 on the plasma membrane expression of clinical CFTR variants. *Cell Chem. Biol.* **30**, 632-642.e5 (2023).
3. Fiedorczuk, K. & Chen, J. Molecular structures reveal synergistic rescue of Δ508 CFTR by Trikafta modulators. *Science* **378**, 284–290 (2022).
4. Bihler, H. *et al.* In vitro modulator responsiveness of 655 CFTR variants found in people with cystic fibrosis. *J. Cyst. Fibros.* S1569199324000213 (2024) doi:10.1016/j.jcf.2024.02.006.
5. Anglès, F., Wang, C. & Balch, W. E. Spatial covariance analysis reveals the residue-by-residue thermodynamic contribution of variation to the CFTR fold. *Commun. Biol.* **5**, 356 (2022).
6. Loureiro, C. A. *et al.* A molecular switch in the scaffold NHERF1 enables misfolded CFTR to evade the peripheral quality control checkpoint. *Sci. Signal.* **8**, ra48–ra48 (2015).
7. Okiyoneda, T. *et al.* Role of calnexin in the ER quality control and productive folding of CFTR; differential effect of calnexin knockout on wild-type and ΔF508 CFTR. *Biochim. Biophys. Acta - Mol. Cell Res.* **1783**, 1585–1594 (2008).
8. Farinha, C. M. & Amaral, M. D. Most F508del-CFTR Is Targeted to Degradation at an Early Folding Checkpoint and Independently of Calnexin. *Mol. Cell. Biol.* **25**, 5242–5252 (2005).
9. Arndt, V., Daniel, C., Nastainczyk, W., Alberti, S. & Höhfeld, J. BAG-2 Acts as an Inhibitor of the Chaperone-associated Ubiquitin Ligase CHIP. *Mol. Biol. Cell* **16**, 5891–5900 (2005).
10. Dai, Q. *et al.* Regulation of the Cytoplasmic Quality Control Protein Degradation Pathway by BAG2. *J. Biol. Chem.* **280**, 38673–38681 (2005).
11. Sondo, E. *et al.* Pharmacological Inhibition of the Ubiquitin Ligase RNF5 Rescues F508del-CFTR in Cystic Fibrosis Airway Epithelia. *Cell Chem. Biol.* **25**, 891-905.e8 (2018).
12. Huang, F., Kirkpatrick, D., Jiang, X., Gygi, S. & Sorkin, A. Differential Regulation of EGF Receptor Internalization and Degradation by Multiubiquitination within the Kinase Domain. *Mol. Cell* **21**, 737–748 (2006).
13. Yu, Y. *et al.* K29-linked ubiquitin signaling regulates proteotoxic stress response and cell cycle. *Nat. Chem. Biol.* **17**, 896–905 (2021).
14. Sumi, T., Matsumoto, K. & Nakamura, T. Specific Activation of LIM kinase 2 via Phosphorylation of Threonine 505 by ROCK, a Rho-dependent Protein Kinase. *J. Biol. Chem.* **276**, 670–676 (2001).
15. Maekawa, M. *et al.* Signaling from Rho to the Actin Cytoskeleton Through Protein Kinases ROCK and LIM-kinase. *Science* **285**, 895–898 (1999).
16. Zhang, Z., Liu, F. & Chen, J. Molecular structure of the ATP-bound, phosphorylated human CFTR. *Proc. Natl. Acad. Sci.* **115**, 12757–12762 (2018).
